# KDM2B promotes cell viability by enhancing DNA damage response in canine hemangiosarcoma

**DOI:** 10.1101/2020.11.17.387704

**Authors:** Kevin Christian M. Gulay, Keisuke Aoshima, Yuki Shibata, Hironobu Yasui, Qin Yan, Atsushi Kobayashi, Takashi Kimura

## Abstract

Epigenetic regulators have been implicated in tumorigenesis of many types of cancer; however, their roles in endothelial cell cancers such as canine hemangiosarcoma (HSA) have not been studied. In this study, we found that lysine-specific demethylase 2B (Kdm2b) was highly expressed in HSA cell lines compared to normal canine endothelial cells. Silencing of Kdm2b in HSA cells resulted to increased cell death *in vitro* compared to the scramble control by inducing apoptosis through the inactivation of the DNA repair pathways and accumulation of DNA damage. Similarly, doxycycline-induced Kdm2b silencing in tumor xenografts resulted to decreased tumor sizes compared to the scramble control. Furthermore, Kdm2b was also highly expressed in clinical cases of HSA, and its expression levels was higher than in hemangioma, a benign counterpart of HSA. Based on these results, we hypothesized that pharmacological Kdm2b inhibition can also induce HSA cell death and can be used as an alternative treatment for HSA. We treated HSA cells with GSK-J4, a histone demethylase inhibitor, and found that GSK-J4 treatment also induced apoptosis and cell death. On top of that, GSK-J4 treatment in HSA tumor-bearing mice decreased tumor sizes without any obvious side-effects. In this study, we demonstrated that Kdm2b acts as an oncogene in HSA by enhancing DNA damage response and can be used as a biomarker to differentiate HSA from hemangioma. Moreover, we indicated that histone demethylase inhibitor GSK-J4 can be used as a therapeutic alternative to doxorubicin for HSA treatment.

## Introduction

Canine hemangiosarcoma (HSA) is a highly malignant tumor of vascular endothelial cells. It is an aggressive tumor with high rates of local recurrence and metastasis, and a low overall survival time^1^. Its high cellular heterogeneity has limited genomic and pathogenesis studies in HSA. Genomic analyses have revealed that HSA cells have somatic coding mutations in the *TP53, PIK3CA*, and *PIK3R1*. Furthermore, *CDKN2A*/B were found to be consistently deleted and copies of *VEGFA, KDR* and *KIT* were gained^2^. The oncogene involved in HSA, however, is still unknown.

Epigenetic mechanisms are essential for reproduction, embryonic development, and maintenance of normal cell function in eukaryotes^3^. Generally, the genetic alterations that cause tumorigenesis are combined with epigenetic shifts, such as aberrant DNA methylation and histone modifications, which may help oncogenic drivers improve cancer development, metastasis, and resistance to therapies^4^. KDM2B (also known as NDY1 and FBXL10), an H3K4me3, H3K36me2/3 and H3K79me3 demethylase, acts as a tumor suppressor in gastric cancer by downregulating glycolysis, and tumor-derived mutation in Kdm2b enhances cell proliferation through the inability of c-FOS degradation^5^. Alternatively, KDM2B can also act as an oncogene in various types of cancers. In leukemia, Kdm2b is highly expressed and is sufficient to transform hematopoietic progenitor cells^6^. In breast cancer, KDM2B regulates polycomb complexes and controls self-renewal of breast cancer stem cells^7^. This bifunctional activity means that the role of KDM2B in tumors is highly context dependent and must be evaluated carefully^8^. While epigenetics is highly involved in pathogenesis of many cancers, its role in HSA is still unknown.

Treatment of HSA is carried out by tumor excision and chemotherapy with doxorubicin, cisplatin, or 5-fluorouracil^9,10^. The mean survival time for surgical treatment is 3 months while the mean survival time for surgical treatment with chemotherapy is less than a year^11,12^. Chemotherapeutic drugs fail to improve survival times of cancer patients due to the non-specificity of their cytotoxic effects in the cardiomyocytes, brain, liver and intestine^13,14^. A more effective, specific, and safer treatment for HSA is warranted.

Histone demethylase inhibitors are effective agents for anticancer treatment. In a previous study, GSK-J4, a histone demethylase inhibitor, decreased AML disease progression by downregulating DNA replication and cell cycle-related pathways through the enrichment of H3K27me3^15^. GSK-J4 was initially developed as a KDM6 inhibitor, but it was found to inhibit the catalytic activity of a wide range of JmjC domain-containing histone demethylases including KDM2B^16^. Epigenetic therapy may provide additional treatment option, but no epigenetic drug has tested for HSA.

In this study, we sought to establish the pathogenesis and to find a therapeutic alternative for HSA by examining the role of epigenetic regulators.

## Results

### Kdm2b is important for HSA cell survival

We first performed RNA-sequencing analysis of canine aortic endothelial cells (CnAOEC) and one HSA cell line, JuB2, to examine the expression levels of epigenetic modifiers. Among the differentially expressed genes, histone methyltransferases/demethylases, histone acetyltransferases/deacetylases, and DNA methyltransferase/demethylases were dysregulated in JuB2 compared to CnAOEC (Fig. 1A, Supplementary Fig. 1A). Next, we performed qRT-PCR to verify RNA-seq results of histone methyltransferases/demethylases in CnAOEC and seven HSA cell lines. Some histone methyltransferases were incoherently expressed in HSA cell lines compared to CnAOEC (Supplementary Fig. 1B). In contrast, three histone demethylases were significantly upregulated in HSA cells compared to CnAOEC. Moreover, one set of paralogs (*KDM2A* and *KDM2B*) were enriched, which implies their importance in HSA (Fig. 1B). We then performed western blotting for Kdm1a, Kdm2a, and Kdm2b and found that their protein levels were upregulated in all HSA cell lines compared to CnAOEC (Fig. 1C and D). To provide further evidence, we performed qRT-PCR analysis in ISO-HAS-B, a human angiosarcoma cell line, compared to Human Umbilical Vein Endothelial Cells (HUVEC). We revealed that histone methyltransferases/demethylases were also dysregulated in ISO-HAS-B (Supplementary Fig. 1C and Fig. 1E). Furthermore, protein expression levels of KDM1A, KDM2A and KDM2B were also upregulated in ISO-HAS-B compared to HUVEC (Fig. 1F and G). These results suggest that epigenetic regulators, specifically Kdm1a, Kdm2a, and Kdm2b, are highly expressed in HSA and human angiosarcoma and, thus, may have a significant role in endothelial cell tumors.

**Fig. 1.**
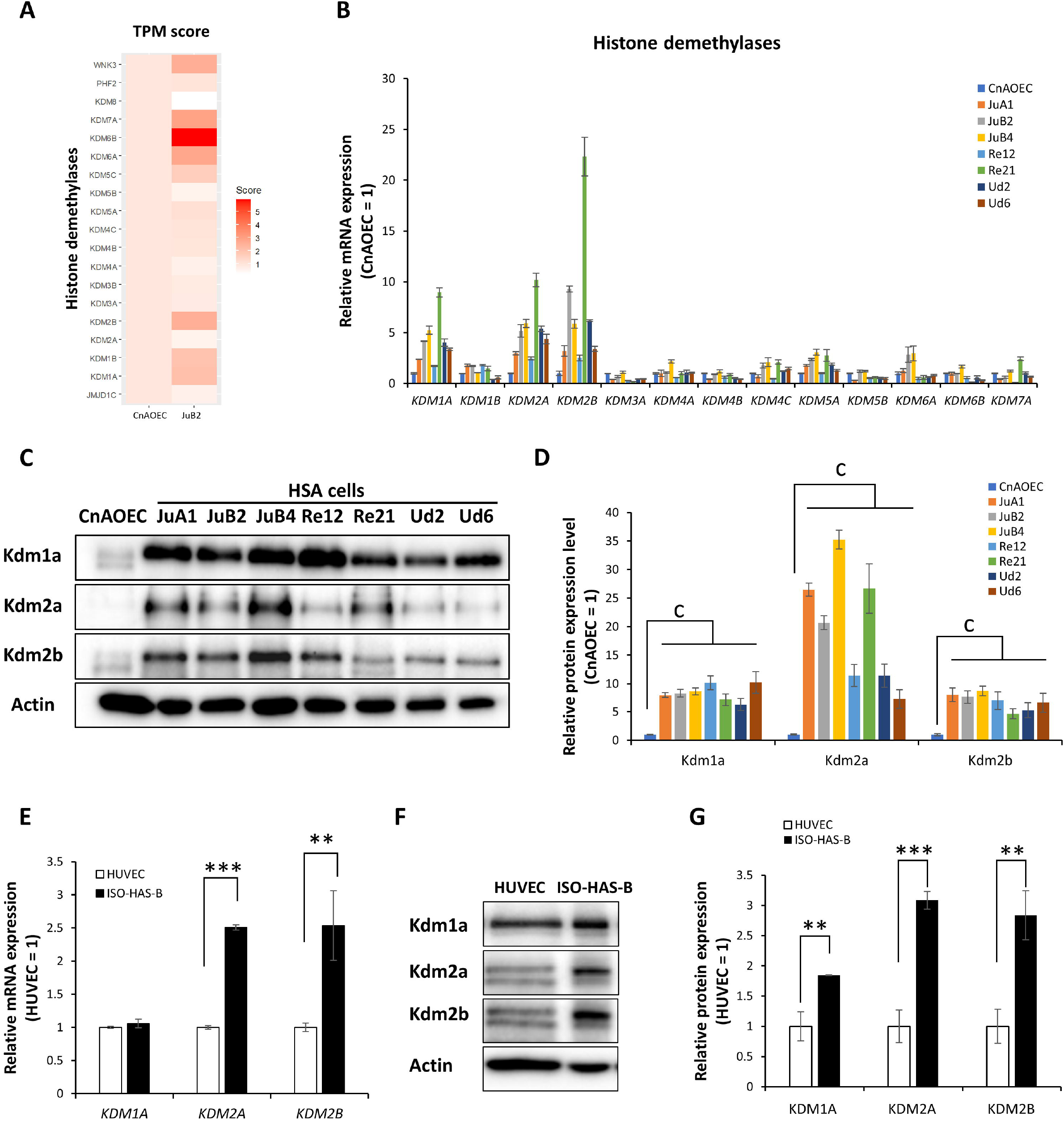
Epigenetic regulators are dysregulated in endothelial cell tumors. **A,** Transcripts per million (TPM) scores of histone demethylases in CnAOEC and JuB2 HSA cell line. **B,** qRT-PCR verification of select histone demethylases in CnAOEC and HSA cell lines. **C,** Western blotting for Kdm1a, Kdm2a, and Kdm2b in CnAOEC and HSA cell lines. **D,** Quantitative analysis of Kdm1a, Kdm2a, and Kdm2b protein expressions in CnAOEC and HSA cell lines. **E,** qRT-PCR analysis of *KDM1A, KDM2A*, and *KDM2B* mRNA expressions in HUVEC and ISO-HAS-B. F, Western blotting for KDM1A, KDM2A, and KDM2B in HUVEC and ISO-HAS-B. **G,** Quantitative analysis of KDM1A, KDM2A, and KDM2B protein expressions in HUVEC and ISO-HAS-B. Data are presented as mean values ± s.d. \**p* <0.05 \*\**p*<0.01 \*\*\**p*<0.001, Student’s *t*-test; ^c^*p*<0.001, Tukey’s test.

To examine whether the overexpression of Kdm1a, Kdm2a and Kdm2b has important functional roles in HSA, we designed three shRNA sequences for each gene to knockdown *KDM1A, KDM2A* and *KDM2B* as well as two scramble RNA controls (scrRNA), and we expressed these shRNAs in HSA cell lines using a doxycycline (Dox)-inducible vector system. Knockdown was verified with western blotting after four days of Dox treatment (Fig. 2A). Knockdown of *KDM1A* and *KDM2A* did not have short term effects in JuB2 viability, whereas knockdown of *KDM2B* markedly decreased JuB2 viability within 4 days (Fig. 2B). To evaluate long term effects of Kdm1a, Kdm2a and Kdm2b on JuB2 cell survival, we performed colony formation assay and found that *KDM1A* and *KDM2A* knockdown could inhibit JuB2 colony formation, whereas *KDM2B* knockdown significantly reduced the number of colonies of JuB2 (Fig. 2C and D, Supplementary Fig. 2A and B). Kdm2b knockdown with constitutively expressed shRNA vectors also induced the same phenotype in JuB2 cells and other HSA cell lines such as JuB4, Re21, and Ud6 (Supplementary Fig. 2C and D). These results suggest that Kdm2b is an important histone demethylase in HSA cell survival, which encouraged us to further investigate its function. To examine whether Kdm2b can alter overall histone methylation levels in HSA, we performed western blotting for histone methylations that can be modified by Kdm2b. However, all possible histone methylations were not affected in JuB2 after *KDM2B* knockdown (Supplementary Fig. 2E). Next, to investigate which Kdm2b domain is responsible for HSA cell viability, we rescued Kdm2b function in *KDM2B* silenced HSA cells. We expressed wild type (WT) canine Kdm2b with silent mutations for the sequences which were targeted by the shRNAs and Kdm2b mutants with a point mutation for the JmjC domain (Kdm2b^H283Y^), the CXXC domain (Kdm2b^C586A^), or with PHD-finger domain deleted (Kdm2b^△PHD^)^17–19^. Overexpression of WT Kdm2b and Kdm2b^C586A^ mutant but not the Kdm2b^H283Y^ or the Kdm2b^△PHD^ mutants rescued the phenotype (Fig. 2E), which suggests that the JmjC and the PHD-finger domains are important for the Kdm2b function in HSA.

**Fig. 2.**
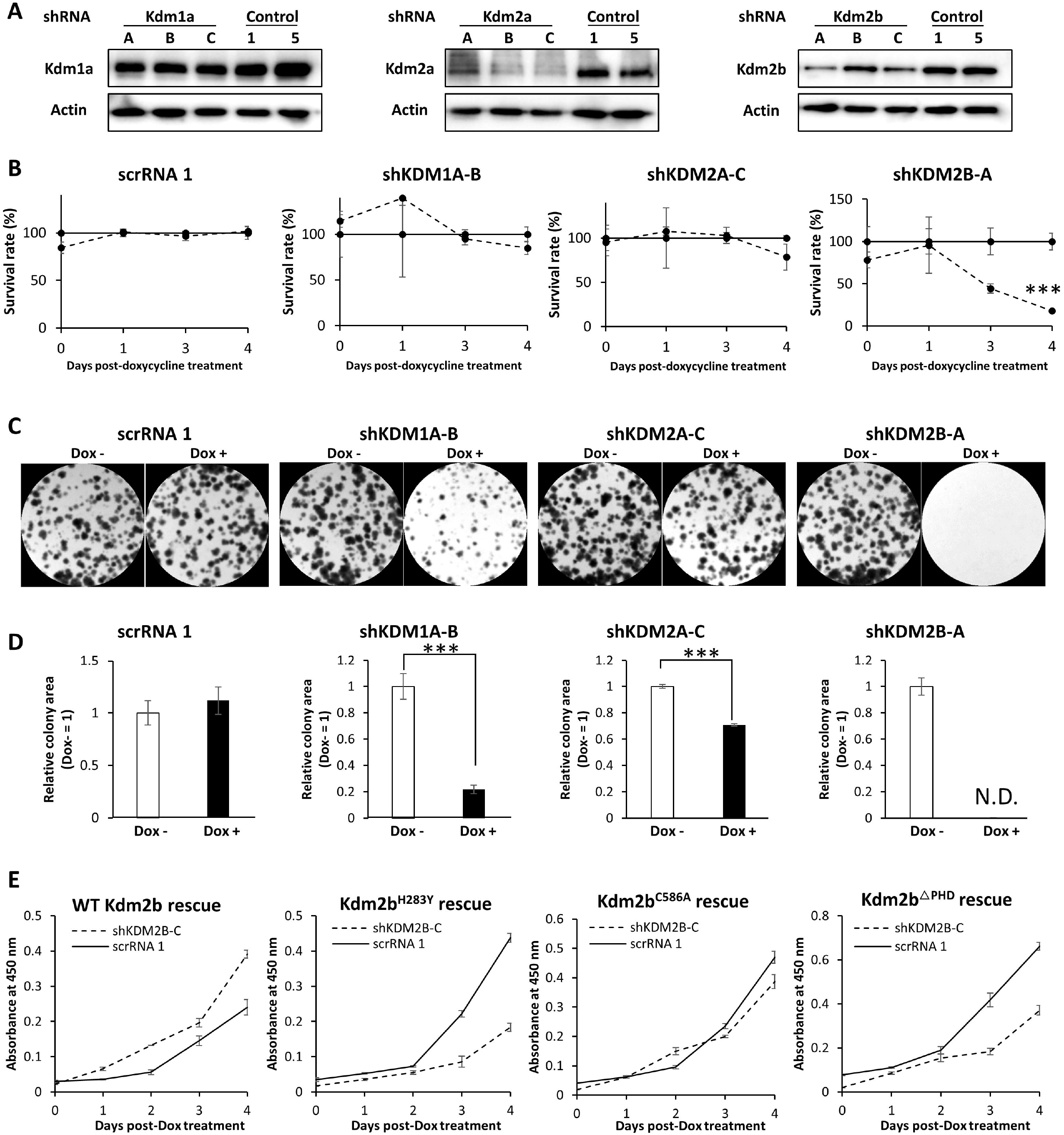
Kdm2b is important for HSA cell survival *in vitro*. **A,** Western blotting of Kdm1a, Kdm2a, or Kdm2b to verify the silencing efficiency of shRNA vectors developed for Kdm1a, Kdm2a, and Kdm2b. **B,** Cell viability analysis after the induction of the inducible shRNA vector using doxycycline. **C,** Colony formation assay of JuB2 cells after Kdm1a, Kdm2a, or Kdm2b silencing. **D,** Quantitative analysis of **C**. **E,** Cell viability analysis after rescue overexpression of WT Kdm2b, or dominant negative mutants for the JmjC (Kdm2b^H283Y^), CxxC (Kdm2b^C586A^), or PHD (Kdm2b^△PHD^) domains. Data are presented as mean values ± s.d. \*\*\**p*<0.001, Two-way ANOVA.

To verify our *in vitro* results and to know the effect of Kdm2b silencing *in vivo*, JuB2 cells with Dox-inducible shRNAs against *KDM2B* or the scramble shRNA were injected subcutaneously into nude mice and the tumor volumes over time were measured. Dox-containing food pellets were provided to the mice to induce shRNA expression when the largest tumor reached 150 mm^3^ in volume. Tumor xenografts started to decrease in volume four days after induction of Kdm2b silencing, moreover, half of the xenografts bearing shKDM2B-A and one out of eight xenografts bearing shKDM2B-C completely regressed 10 days post-induction (Fig 3A and B). At the endpoint, tumor xenografts with silenced Kdm2b were significantly smaller compared with the xenografts overexpressing the scramble RNA control (Fig. 3C). We also xenografted JuB2 cells expressing the silencing vector for Kdm1a or Kdm2a. Although tumor volume repression was initially observed post-doxycycline treatment, tumor regrowth was observed 10 post-doxycycline treatment signifying that other histone demethylases or proteins can rescue Kdm1a or Kdm2a function *in vivo* or some cell populations were not affected by their knockdown (Fig. 3A).

**Fig. 3.**
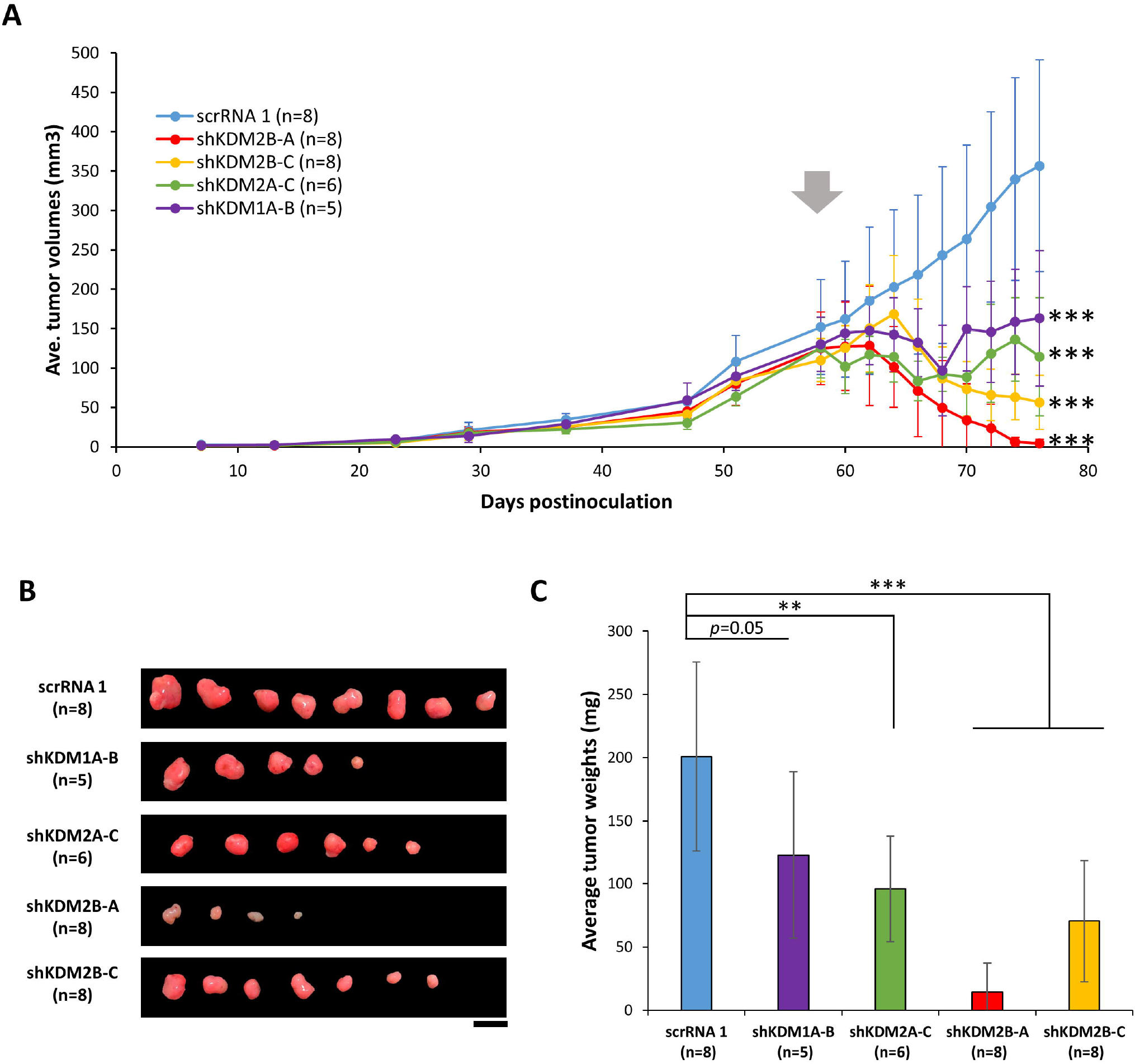
Kdm2b is important in HSA cell survival *in vivo*. **A,** Tumor growth at different time points before and after the induction of the shRNA expression in JuB2 cells inoculated in nude mice. \*\*\**p*<0.001, two-way ANOVA with Dunnett’s post-hoc test for tumor volumes after starting doxycycline treatments (arrow). **B and C,** Tumor sizes and weights of tumors harboring scramble RNA or shRNA for Kdm1a, Kdm2a, or Kdm2b 78 days after tumor transplantation. Scale = 1 cm. \*\**p*<0.01 \*\*\**p*<0.001, Dunnett’s test. Data are presented as mean values ± s.d.

These results suggest that Kdm2b is important for HSA cell survival both *in vitro* and *in vivo* and its local catalytic activity is responsible for the phenotype.

### *KDM2B* knockdown induces apoptosis via accumulation of DNA damages

To understand the mechanisms by which Kdm2b regulates HSA cell survival, we performed RNA-seq followed by gene set enrichment analyses (GSEA) in JuB2 cells expressing shKDM2B-A versus JuB2 cells expressing scrRNA-1. Silencing of Kdm2b showed negative correlations with angiogenesis and glycolysis pathways, which might reflect decreased HSA cell viability (Supplementary Fig. 3A). Interestingly, expressions of genes downregulated in response to ultraviolet (UV) radiation (HALLMARK_UV_RESPONSE_DN) were decreased by Kdm2b knockdown (Fig. 4A). Considering this correlation and the fact that UV response can trigger DNA damage response^20^, we speculated that DNA damage might be related to *KDM2B* knockdown phenotypes. This speculation was further supported by positive enrichment of the G2M checkpoint pathway and the interferon response pathway, which has been reported to be associated with mitochondrial and nuclear DNA damage (Supplementary Fig. 3A)^21–24^. These results were validated by qRT-PCR (Fig. 4B, Supplementary Fig. 3B to E). To further dissect the mechanism, we examined the expression levels of proteins involved in DNA repair pathway. Total Atm, c-Fos, and γH2A.X expressions were upregulated in HSA cell lines compared to CnAOEC and their expressions were significantly decreased by *KDM2B* knockdown (Fig 4C and D). Phosphorylated-Atm (pAtm), an active form of Atm, was also decreased by *KDM2B* knockdown. Next, we carried out alkaline comet assay to detect DNA strand breaks in HSA cell lines upon *KDM2B* silencing. Our results showed that *KDM2B* silencing drastically increased tail DNA percentages, tail lengths, and tail momentums compared to the scramble control (Fig. 4E and F). Furthermore, flow cytometry analysis using propidium iodide (PI) revealed that JuB2 with silenced *KDM2B* increased aneuploid peaks although parental HSA cells also showed aneuploidy (Fig 5A and B, Supplementary Fig. 4A and B). In contrast, overexpression of WT Kdm2b in JuB2 HSA cells resulted to significantly decreased aneuploid cell population compared to empty vector (EV) expressing JuB2 cells (Fig 5C and D). In addition, active apoptosis markers, cleaved-caspase 3, Bax and phosphorylated Erk1/2, were increased in cells with silenced *KDM2B* compared to the scramble control cells (Fig 5E). Furthermore, CnAOEC overexpressing WT Kdm2b continued proliferating beyond day 20 while the EV expressing CnAOEC showed decreased cell proliferation and cell death starting at day 11 (Fig 5F). In addition, Kdm2b overexpression decreased the phosphorylation of Erk1/2 proteins at day 19 post-lentiviral Kdm2b overexpression (Fig. 5G and H) and repressed mRNA expressions of cell cycle checkpoint genes such as *p15^ink4b^, p16^ink4a^*, and *ATR* (Fig. 5I). These results suggest that *KDM2B* knockdown in HSA cells induces cell death via apoptosis caused by accumulation of DNA damages due to low expression of proteins involved in DNA repair.

**Fig. 4.**
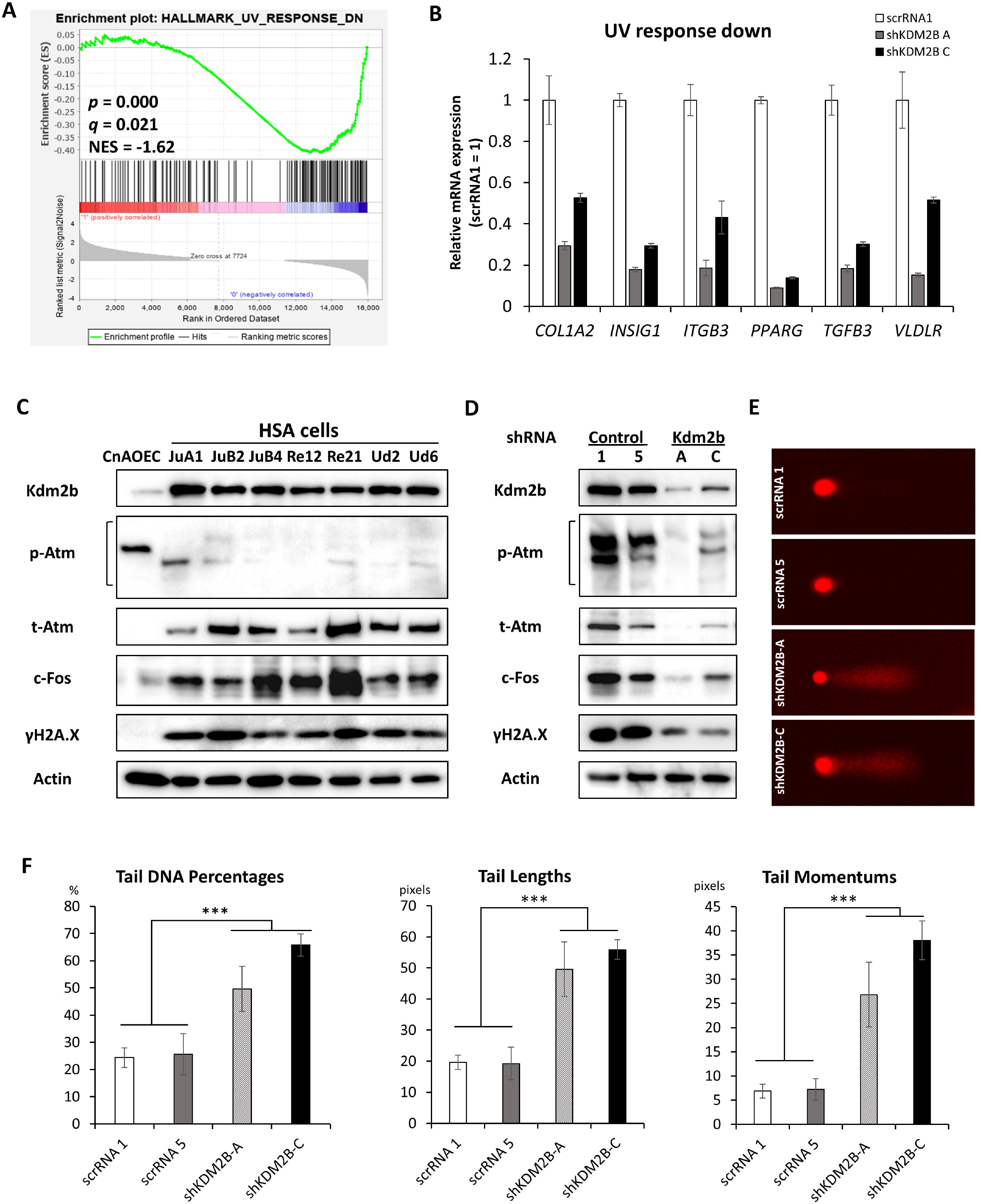
Kdm2b positively regulates DNA repair pathway. **A,** GSEA enrichment plot for UV_Response_DN in shKDM2B-A versus scrRNA-1. **B,** qRT-PCR verification of gene expressions listed in UV_Response_DN gene set. **C and D,** Western blotting for Atm, c-Fos, and γH2A.X in normal endothelial cells and HSA cell lines (**C**) and in JuB2 cells expressing shRNA control or shKDM2B (**D**). **E,** Representative images of alkaline comet assays in scrRNA or shKDM2B JuB2 cells. F, Tail DNA percentages, lengths, and momentums of JuB2 HSA cells harboring scrRNA or shRNA for Kdm2b. Data are presented as mean values ± s.d. \*\*\**p*<0.001, Tukey’s test.

**Fig. 5.**
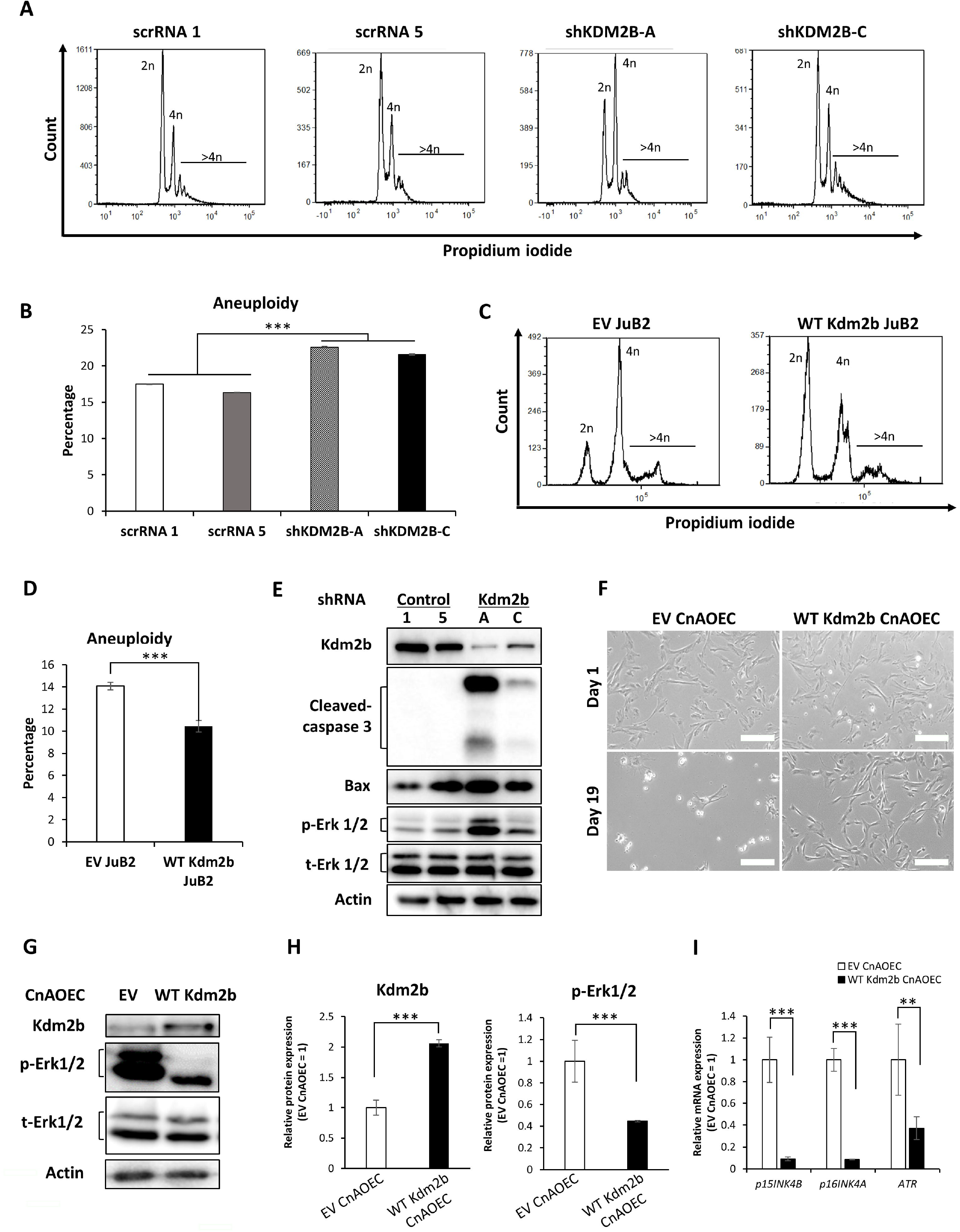
Kdm2b regulates aneuploidy and apoptosis in HSA. **A,** Histograms of PI intensities in scrRNA and shKDM2B JuB2 cells. **B,** Percentages of aneuploid cells in shRNA and shKDM2B JuB2 cells. \*\*\**p*<0.001, Tukey’s test. **C,** Histograms of PI intensities in EV infected JuB2 cells and WT Kdm2b JuB2 cells. **D,** Percentages of aneuploid cells in EV JuB2 and WT Kdm2b JuB2. \*\*\**p*<0.001, Student-*t* test. **E,** Western blotting for cleaved-caspase 3, Bax and Erk 1/2 in JuB2 HSA cells expressing scrRNA controls or shKDM2B RNAs. F, Phase contrast images of CnAOEC cells infected with EV or overexpressing WT Kdm2b at day 1 and day 19. Scale = 100 μm. **G,** Western blotting of Erk 1/2 in CnAOEC infected with EV or overexpressing WT Kdm2b. **H,** Quantification of Kdm2b and phosphorylated Erk 1/2 expression in EV infected or WT Kdm2b expressing CnAOEC normalized with Actin and total Erk 1/2, respectively. **I,** qRT-PCR analysis of cell cycle related genes after overexpression of WT Kdm2B in normal endothelial cells. Data are presented as mean values ± s.d. \*\**p*<0.01 \*\*\**p*<0.001, Student’s *t*-test.

### Kdm2b is highly expressed in clinical cases of HSA

We demonstrated that Kdm2b upregulation was important for HSA cell survival, but Kdm2b expression in clinical HSA cases has not been investigated. To this end, we analyzed seventeen clinical cases of HSA and compared the immunohistochemical Kdm2b expression between HSA cells and normal endothelial cells in the same sample. Our results showed that Kdm2b was highly expressed in HSA cells compared to normal endothelial cells in almost all the cases examined regardless of proliferation patterns or degree of differentiation (Fig 6A and B). The average Kdm2b score in HSA cells was significantly higher than that in normal endothelial cells (Fig. 6C). We then also examined the benign endothelial tumor, hemangioma (HMA), for Kdm2b expression. Interestingly, Kdm2b expression levels in hemangioma cells were likely lower than normal endothelial cells in the same sample (Fig. 6D to F, *p*=0.096). The average Kdm2b expression scores in HSA were significantly higher than in HMA cells when the average Kdm2b scores in tumor cells were normalized to normal endothelial cells in the same slides (Fig. 6G). Kdm2b expressions were also analyzed in histologically-similar tumors, hemangiopericytoma (HPC), melanoma, and fibrosarcoma cells. Results showed that HPC, melanoma, and fibrosarcoma cells had similar Kdm2b expression compared to normal endothelial cells in the same slide (Supplementary Fig. 5A to C). These results suggest that Kdm2b expresses in clinical HSA samples and can be used as a biomarker to differentiate HSA and HMA. In addition, Kdm2b may also be used as a molecular marker to differentiate HSA from HPC, melanoma, and fibrosarcoma.

**Fig. 6.**
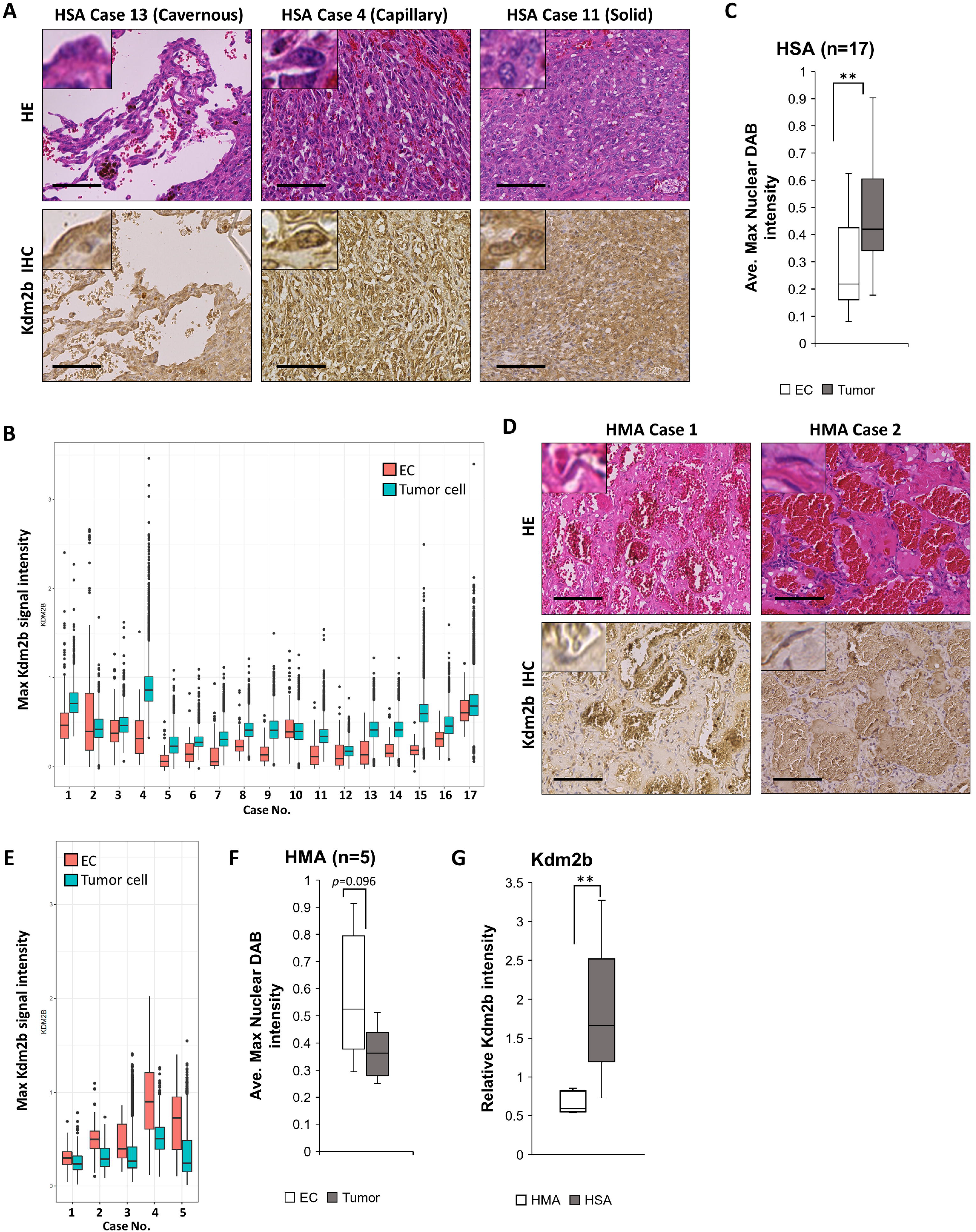
Kdm2b can be used as a differential biomarker for HSA. **A,** Representative images of hematoxylin-eosin staining (HE) and immunohistochemistry (IHC) for clinical HSA samples. Proliferation patterns are indicated with case numbers. Scale = 100 μm. **B,** Max nuclear DAB intensities in individual endothelial cells (EC) and tumor cells (Tumor) of each HSA case. **C,** Average max nuclear DAB intensities of EC and tumor cells in each HSA case. **D,** Representative HE and IHC images of hemangioma (HMA), a benign endothelial cell tumor. Scale = 100 μm. **E,** Max nuclear DAB intensities in individual EC and tumor cells of each HMA case. F, Average max nuclear DAB intensities of EC and tumor cells in each HMA case. **G,** Comparison of relative Kdm2b intensity of HSA and HMA cells normalized by normal endothelial cells in their respective slides. Data are presented as mean values ± s.d. \*\**p*<0.01, Student’s *t*-test.

### GSK-J4 inhibits HSA cell viability

We demonstrated that KDM2B inhibition through shRNA could induce HSA cell death. Based on this, we hypothesized that pharmacological Kdm2b inhibition could also induce HSA cell death. To test this hypothesis, we treated HSA cell lines with different concentrations of GSK-J4 and compared it to doxorubicin-treated cells. GSK-J4 inhibited HSA cell viability at a lower concentration in JuB2, JuB4, and Ud6 cell lines compared to doxorubicin (Fig. 7A). HSA cell lines treated with GSK-J4 have higher expressions of cleaved-caspase 3 compared to HSA cells treated with doxorubicin. In contrast, GSK-J4 treated HSA cells have decreased t-Atm and γH2A.X expression compared to HSA cells treated with DMSO control (Fig. 7B).

**Fig. 7.**
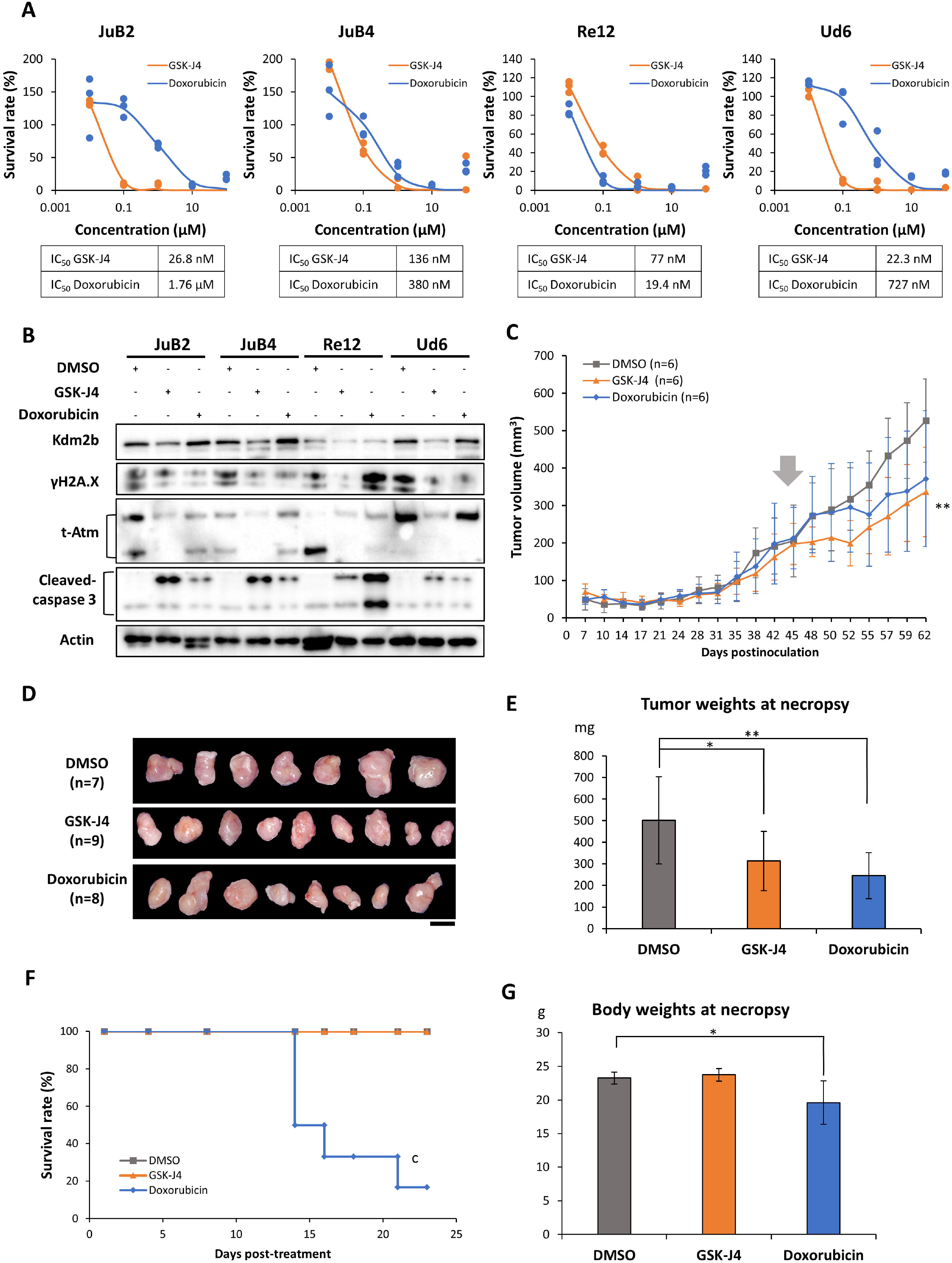
GSK-J4 inhibits HSA cell proliferation *in vitro* and *in vivo*. **A,** Survival rates and IC_50_ values of GSK-J4 or doxorubicin-treated HSA cell lines. **B,** Western blotting for γH2A.X, Atm and cleaved-caspase 3 in HSA cells treated with GSK-J4 or doxorubicin. **C,** Tumor growth curves of JuB2 HSA cell xenografts in nude mice. Treatment with DMSO, GSK-J4, or doxorubicin started at day 45 (arrow). n indicates the number of mice. \*\**p* < 0.01, two-way ANOVA with Dunnett’s post-hoc test. **D,** Gross images of collected tumors at day 64. n indicates the number of collected tumors. **E,** Tumor weights of DMSO, GSK-J4, or doxorubicin-treated nude mice at necropsy. F, Kaplan-Meier survival curves of DMSO, GSK-J4, or doxorubicin-treated nude mice after starting treatments. \*\**p* < 0.01, Log-rank test. **G,** Body weights of DMSO, GSK-J4, or doxorubicin-treated nude mice at necropsy. Data are presented as mean values ± s.d. \*\**p*<0.01, Tukey’s test.

To determine whether GSK-J4 can be used as an alternative drug for HSA treatment, we treated nude mice harboring JuB2 tumors with DMSO, GSK-J4 or doxorubicin. GSK-J4 treatment significantly decreased the tumor growth over time and tumor weights at the endpoint compared to DMSO treatment (Fig 7C-E). Although Doxorubicin treatment also led to decreased tumor growth, it had significant toxicity and led to 83% mortality in 21 days (Fig. 7F). In contrast, GSK-J4 did not induce any death during the treatment period. Body and liver weights decreased in doxorubicin-treated mice compared to DMSO or GSK-J4 treated mice (Fig. 7G, Supplementary Fig. 6A). In addition, increased hematopoiesis in bone marrow, a sign of myelosuppression, and dilation of intestines were only observed in mice treated with doxorubicin (Supplementary Fig. 6B and C). These results suggest that GSK-J4 can selectively inhibit HSA cell growth by inducing HSA cell apoptosis without any obvious side-effects. Thus, GSK-J4 could be used as a therapeutic alternative to doxorubicin for HSA treatment.

## Discussion

To our knowledge this is the first study which evaluated the role of epigenetic regulators in HSA. We demonstrated that Kdm2b upregulation in HSA cell lines and clinical samples is important for HSA cell survival by regulating DNA damage repair and apoptosis. There are three possible reasons for the high expression of Kdm2b in HSA. Firstly, *KDM2B* upstream regulators may be dysregulated. In bladder cancer, the upregulation of fibroblast growth factor-2 upregulated the KDM2B/EZH2-miR-101 pathway and promoted tumor cell proliferation, survival, and migration^25^. In squamous cell and cervical carcinomas, increased copy number of *MYC* resulted to increased KDM2B expression^26^. Secondly, the *KDM2B* gene itself may have acquired increased copy numbers. We demonstrated that parental HSA cell lines contain an aneuploid cell population. Since cell aneuploidy can cause genomic instability as a result of decreased DNA damage repair activity^27^, some genes including *KDM2B* in HSA might have increased copy numbers. Lastly, *KDM2B* might be upregulated to compensate for DNA damage and to integrate genetic stability in HSA genomes. As we mentioned above, flow cytometry analysis revealed multiple aneuploid peaks in HSA cell lines. Aneuploidy has also been reported in clinical cases of HSA and in HSA cell lines, which implies that HSA cells are subject to high level cellular stress^28–30^. In such conditions, HSA cells may bypass cell death caused by accumulated cellular stresses through modulation of gene expression. We showed that knockdown of Kdm2b in HSA cell lines decreased DNA repair protein expressions and increased DNA damage. In contrast, exogenous expression of Kdm2b in JuB2 and in CnAOEC increased euploid population and inhibited apoptosis, respectively. Kdm2b may ease cellular stress caused by DNA damage in endothelial cells by regulating the DNA repair system.

KDM2B and other histone demethylases are known to have a double-edged function in cancer^8,31^. A study showed that KDM2B suppresses tumorigenesis in gastric cancer by enhancing c-Fos degradation, and that impairment of KDM2B through patient-derived mutations enhances tumor cell proliferation^32^. In contrast, several studies showed that degradation of c-FOS by KDM2B in prostate cancer and glioblastoma multiforme can increase cancer cell resistance to chemotherapy^33,34^. In HSA, we showed that positive regulation of these genes by Kdm2b can enhance HSA cellular viability in contrast to a previous report where KDM2B was shown to promote colon cancer cell survival by negatively regulating DNA damage response-related genes such as ATM and ATR^35^. These contrasting evidences support the role of KDM2B in DNA damage response and suggest that the role of KDM2B in DNA damage response in cancer cells is highly context dependent.

We have presented various evidences which demonstrate the role of Kdm2b as an oncogene in HSA by regulating DNA repair system and apoptosis, and that Kdm2b can be used as a biomarker to aid HSA diagnosis. We also demonstrated that a histone demethylase inhibitor GSK-J4 can work as a new therapeutic alternative to doxorubicin treatment in HSA. These findings shed light on the epigenetic pathology and provides a new insight for a novel therapy in HSA.

## Materials and Methods

### Cell culture

The HSA cell lines were donated by Dr. Sakai (Gifu University)^36^ and cultured as described previously^37^. CnAOEC was purchased from Cell Applications (CA, USA), and ISO-HAS-B was received from Cell Resource Center for Biomedical Research Cell Bank in Tohoku University (Sendai, Japan)^38^. Both CnAOEC and ISO-HAS-B were cultured in Endothelial Cell Growth Medium 2 Kit (Takara Bio, Inc. Kusatsu, Japan). All cells used were routinely tested for *Mycoplasma* using PCR and were submitted to ICLAS Monitoring Center (Kawasaki, Japan) for Mouse hepatitis virus testing^39,40^.

### Mice

All mouse experiments were performed under the AAALAC guidelines in Yale University (protocol number: 2018-11286) and Hokkaido University (protocol number:19-0130). Seven-week old female Balb/c Nude mice purchased from Charles River Laboratories (MA, USA) were used for *KDM2B* knockdown experiments. Six-week-old KSN/Slc mice purchased from Japan SLC, Inc. (Shizuoka, Japan) were used for drug treatment experiments. Mice were kept in a temperature-controlled specific-pathogen-free facility on a 12 hr light/dark cycle. Animals in all experimental groups were examined at least twice weekly for tumorigenesis.

### Tumor xenograft studies

A day before tumor inoculation, KSN/Slc mice were treated with 100 μL of 2.5 mg/mL anti-asialo GM1 (Fujifilm Wako Pure Chemical Industries, Osaka, Japan) to increase the success rate of transplantation by depleting NK cells^41^. JuB2 parental HSA cells and JuB2 cells expressing the shRNAs or the scramble shRNA for *KDM2B* in the presence of doxycycline were cultured in 15 cm dishes without doxycycline. Mice were randomly assigned in each group. Three million HSA cells were resuspended in Corning^®^ Matrigel^®^ Basement Membrane Matrix (Corning Inc. NY, USA) and inoculated subcutaneously in mice anesthetized with 3% isoflurane or 100 mg/kg Ketamine and 10 mg/kg Xylazine. Tumor sizes were measured twice weekly one week after inoculation. When the largest tumor reached 150 mm^3^ in volume, mice were fed doxycycline-containing food to induce shRNA expression, or treated thrice weekly for three weeks with 50 mg/kg DMSO, 50 mg/kg GSK-J4 (Medchemexpress, NJ, USA), or 5 mg/kg doxorubicin (Fujifilm Wako Pure Chemical Industries) intraperitoneally. Mice were euthanized with CO_2_ when tumors reached 500 mm^3^ in volume or when mice exhibited abnormal behavior. Tumors and major organs were weighed and fixed in 10% neutral buffered formalin for histological examination.

### Western Blotting

SDS lysis buffer (2% SDS, 50 mM Tris-HCl (pH6.8), 1mM EDTA (pH 8.0)) with EDTA-free proteinase inhibitor cocktail (Sigma-Aldrich, MO, USA) was added in cultured cells. Cell lysates were then sonicated using BRANSON Sonifier 450 (Branson Ultrasonics Corporation, CT, USA) for two secs. Protein concentrations were measured with the Pierce™ BCA Protein Assay Kit (Thermo Fisher Scientific) before adding 4X Sample loading buffer (200 mM Tris-HCl buffer (pH 6.8), 8% SDS, 40% Glycerol, 1% bromophenol blue, 20% 2-mercaptoethanol) and denaturing at 98°C for 10 mins. 3 μg proteins were separated by SDS-PAGE and electrotransferred onto Immobilon^®^-P transfer membranes (Merck Millipore, MA, USA), blocked with 5% skim milk in Tris-buffered saline with 5% Tween 20 (TBST), or 5% BSA in TBST for one hour at room temperature (RT) and incubated with primary antibody in Can Get Signal Solution^®^ 1 (TOYOBO, Osaka, Japan) overnight at 4°C. The membranes were washed with TBST three times before incubating with the corresponding secondary anti-mouse or anti-rabbit IgG antibody (GE Healthcare) in Can Get Signal Solution^®^ 2 (TOYOBO). Signals were developed with Immobilon^®^ Western Chemiluminescent HRP substrate (Merck Millipore) and visualized in ImageQuant LAS 4000 mini luminescent image analyzer (GE Healthcare). Captured data were processed using ImageJ software^42^. The list of antibodies used in this study can be found as Supplementary Table 1.

### Quantitative RT-PCR (qRT-PCR)

Total RNA was extracted with Nucleospin^®^ RNA isolation kit (Macherey-Nagel GmbH & Co. Düren, Germany) following the manufacturer’s instructions. Synthesis of cDNA was performed using the PrimeScript™ Reverse Transcriptase (Takara Bio, Inc.) according to the manufacturer’s instructions. qRT-PCR was performed with StepOne™ Real Time System (Thermo Fisher Scientific). The oligos for qRT-PCR were designed as described elsewhere^37,43^, and listed within Supplementary Table 2.

### RNA-sequencing

CnAOEC or JuB2 HSA cells were cultured in 10 cm dishes in triplicate. Upon reaching 80% confluency, RNA was extracted with Nucleospin^®^ RNA isolation kit (Macherey-Nagel GmbH & Co.) following the manufacturer’s instructions. RNA samples were submitted to Annoroad (Beijing, China) for further analyses. Quality testing was carried by measuring RNA integrity (RIN), OD_260/280_ and OD_260/230_. All samples had an RIN of 9.3 or better and OD readings were within the range of 1.8-2.2. RNA-seq libraries were constructed using NEBNext^®^ Ultra RNA Library Prep Kit for llumina^®^ (New England Biolabs, MA, USA) and sequenced with the Illumina HiSeq X-Ten platform (Illumina, CA, USA) to generate a minimum of 20 million paired-end 150 bp reads. Sequencing reads were mapped to the canine reference genome CanFam3.1 using STAR and aligned using RSEM^44,45^. Differential expression analyses were carried out using edgeR^46^, and gene expression profiles were analyzed by GSEA v4.03^47,48^. The gene set database of h. all. v7.1. symbols. gmt (Hallmarks) was used.

### shRNA vector construction

The shRNAs used in this study were designed using the Hannon lab shRNA design tool (http://hannonlab.cshl.edu/GH_shRNA.html, Cold Spring Harbor Laboratory). Oligos were inserted to pLKO.1-TRC or pInducer10-mir-RUP-PheS vector, a gift from David Root (Addgene plasmid # 10878; http://n2t.net/addgene: 10878; RRID:Addgene_10878)^49^ and from Stephen Elledge (Addgene plasmid # 44011; http://n2t.net/addgene:44011; RRID:Addgene 44011)^50^, respectively. shRNA expressions were induced by supplementing 2 μM doxycycline in the culture medium. The list of oligonucleotide sequences used for shRNA knockdown can be found as Supplementary Table 3.

### Overexpression vector construction

The coding sequence of canine *KDM2B* (ENSCAFT00000093772.1) was cloned from cDNA synthesized from canine heart. Mutant Kdm2b (H283Y, C586A, ΔPHD) were synthesized as described elsewhere^15,51^. The amplicon tagged with FLAG sequences at its 3’ end was ligated into CSII-CMV-MCS-IRES2-Bsd vector, a gift from Dr. Miyoshi (RIKEN BioResource Center, Ibaraki, Japan), with In-Fusion^®^ HD Cloning Kit (Takara Bio, Inc.) according to the manufacturer’s instruction. The list of oligonucleotide sequences used for CDS cloning are can be found as Supplementary Table 4.

### Lentivirus production

Lentiviruses were produced following a protocol described elsewhere with a slight modification that virus containing culture medium was used without concentration^37^. Selection of positive clones were done by culturing of cells in 10 μg/ml blasticidin- or 4 μg/ml puromycin-containing cell medium.

### Alkaline comet assay

JuB2 overexpressing shRNA for *KDM2B* or scrRNA were cultured in doxycycline-containing medium for four days. 4×10^4^ cells were used for alkaline comet assay as described previously. 20 μg/mL PI was used for staining and the comets were visualized with BZ-9000 (BIOREVO) fluorescence microscope (Keyence, Osaka, Japan). Experiments were performed at least three times with triplicates.

### Cell cycle analysis

When HSA cells reach 70% confluency, they were stained with 1 μL of 0.1M BrdU for 45 mins at 37°C. Cells were washed with phosphate-buffered saline (PBS) and trypsinized routinely. The cells were fixed in 70% ethanol overnight and washed with 0.5% Triton X-100 in PBS (PBST) before resuspending the cells in 500 μL of 2N HCl-0.5% Triton X-100 for 30 mins at RT and neutralizing with 500 μL of 0.1M Na_2_B_4_O_7_·10H_2_O (pH 8.5) for 30 mins at RT. After blocking with 1% BSA-0.3% Triton X-100 in PBS for 1 hr and washing with PBST, cells were counted and divided into two tubes; 2.5X10^5^ cells were used as controls and incubated in the blocking buffer while 6X10^5^ cells were incubated with anti-BrdU monoclonal antibody (1:100; MOBU-1 clone, B35128, Thermo Fisher Scientific) for 1 hr at RT. Excess primary antibodies were washed before staining with AlexaFluor 488 (1:1000; Thermo Fisher Scientific). DNA was stained with 10 μg/200 μL Propidum Iodide (Dojindo Molecular Technologies, Inc., Kumamoto, Japan). Cell cycle and cell proliferation were analyzed in BD FACSVerse™ flow cytometer (BD Biosciences, NJ, USA). Results were analyzed with FCS Express 4 software (De Novo Software, CA, USA). Experiments were performed at least three times with triplicates.

### Cell viability analysis

Cell viability was measured with Cell Counting Kit-8 (Dojindo Molecular Technologies, Inc.) according to the manufacturer’s instructions. The absorbance at 450 nm was measured with NanoDrop™ 2000 (Thermo Fisher Scientific). Determination of IC_50_ were performed using Ky Plot 6.0 software (KyensLab, Inc., Tokyo, Japan) as described elsewhere^52^. Experiments were performed at least three times with triplicates.

### Colony formation assay

500 HSA cells were seeded in 6-well culture plates containing 2 mL normal medium supplemented with DMSO or 2 μM doxycycline. Cells were cultured until the largest colony reached 2 mm in diameter. Cells were fixed with 4% paraformaldehyde for 20 mins at RT, stained with 0.01% Crystal Violet (Sigma-Aldrich, MO, USA) for 30 mins at RT, and the images were captured with ChemiDoc XRS Plus (Bio-rad, CA, USA). Colony areas were measured using ColonyArea plugin for ImageJ^42,53^. Experiments were performed at least three times with triplicates.

### Histopathology and immunohistochemistry (IHC)

Written informed consents were obtained from the owners of the patient dogs, and the samples were used only for research purposes. Hematoxylin and eosin staining was performed as described previously^54^. For IHC, tissue sections were deparaffinized, and heat-induced antigen retrieval was performed in citric acid buffer (pH 6.8) with a pressure cooker. Endogenous peroxidases were quenched with 0.3% H_2_O_2_ in methanol for 15 min at RT before blocking the tissue sections with 10% normal rabbit serum (Nichirei biosciences, Tokyo, Japan) for an hour at RT and incubating with KDM2B antibody (sc-293279, 1:50, Santa Cruz Biotechnology, CA, USA) overnight at 4°C. Slides were washed with 0.01M PBS before incubating with rabbit anti-mouse antibody (Nichirei biosciences) for 30 min at RT. Signals were developed with 3,3’-diaminobenzidine tetrahydrochloride (DAB; Dojindo).

### Quantification of IHC scores

Histological slides were scanned with Nano Zoomer 2.0-RS (Hamamatsu Photonics, Hamamatsu, Japan) and processed in QuPath ver 0.2.1^55^. Scanned slides were opened as Brightfield (H-DAB) in QuPath, and staining colors were automatically adjusted using Estimate Stain Vectors function. Healthy tumor regions or normal blood vessels were randomly selected, and cells were detected using Cell Detection function. Cells were annotated based on their morphologies and location to allow QuPath to automatically classify each cell type correctly using the Create Detection Classifiers function. The measured data was exported and used for further analysis. We used Nuclear: DAB OD max scores for quantitative analysis because Kdm2b signals were found only in heterochromatin regions in nuclei but they had abundant euchromatin region. Furthermore, nonspecific signals from fibrin and erythrocytes were often misinterpreted as nuclear or cytoplasmic signals especially in normal endothelial cells, thus, to make the analysis as precise as possible, we decided to use max intensities in the nuclei.

### Statistical analysis

Statistical analyses were performed with Microsoft Excel and R software (version 3.6.3). Student’s *t*-tests was used to analyze difference between two groups while the Tukey’s test was used to analyze differences between multiple groups. The significance of the differences in tumor volumes of DMSO, GSK-J4 or doxorubicin-treated mice were compared using Dunnet’s test while their survival post-treatment were compared using Log-rank test. *p*-values less than 0.05 were considered statistically significant.

## Acknowledgements

We are grateful to Dr. Osamu Ichii, Dr. Junpei Yamazaki, and Dr. Noboru Sasaki for their invaluable support during the conduct of the study. We appreciate useful discussions with the members of the Laboratory of Comparative Pathology, Faculty of Veterinary Medicine, Hokkaido University, and the members of Yan laboratory, Department of Pathology, Yale School of Medicine. This research was supported by the Sasagawa Scientific Research Grant for Young Researchers (KG, Research No. 2019-4111) provided by the Japan Science Society and the KAKENHI Grant-in-Aid for Young Scientist (KA, Number 18K14575 and 20K15654) provided by Japan Society for the Promotion of Science.

## Availability of data and materials

The RNA sequence data will be deposited in NCBI’s Gene Expression Omnibus. All other data supporting the findings of this study can be found within the supplementary files.

## Author contributions

KG and KA designed the experiments. KG performed in vitro experiments. KG and KA performed in vivo experiments. YS and HY assisted with the comet assay. KA analyzed the RNA-sequencing data and performed other bioinformatics analyses. KG and KA were responsible for the statistical analyses. QY, AK and TK supervised the experiments. KG and KA wrote the paper.

