## Supplementary data for "KDM2B promotes cell viability by enhancing DNA damage response in canine hemangiosarcoma"

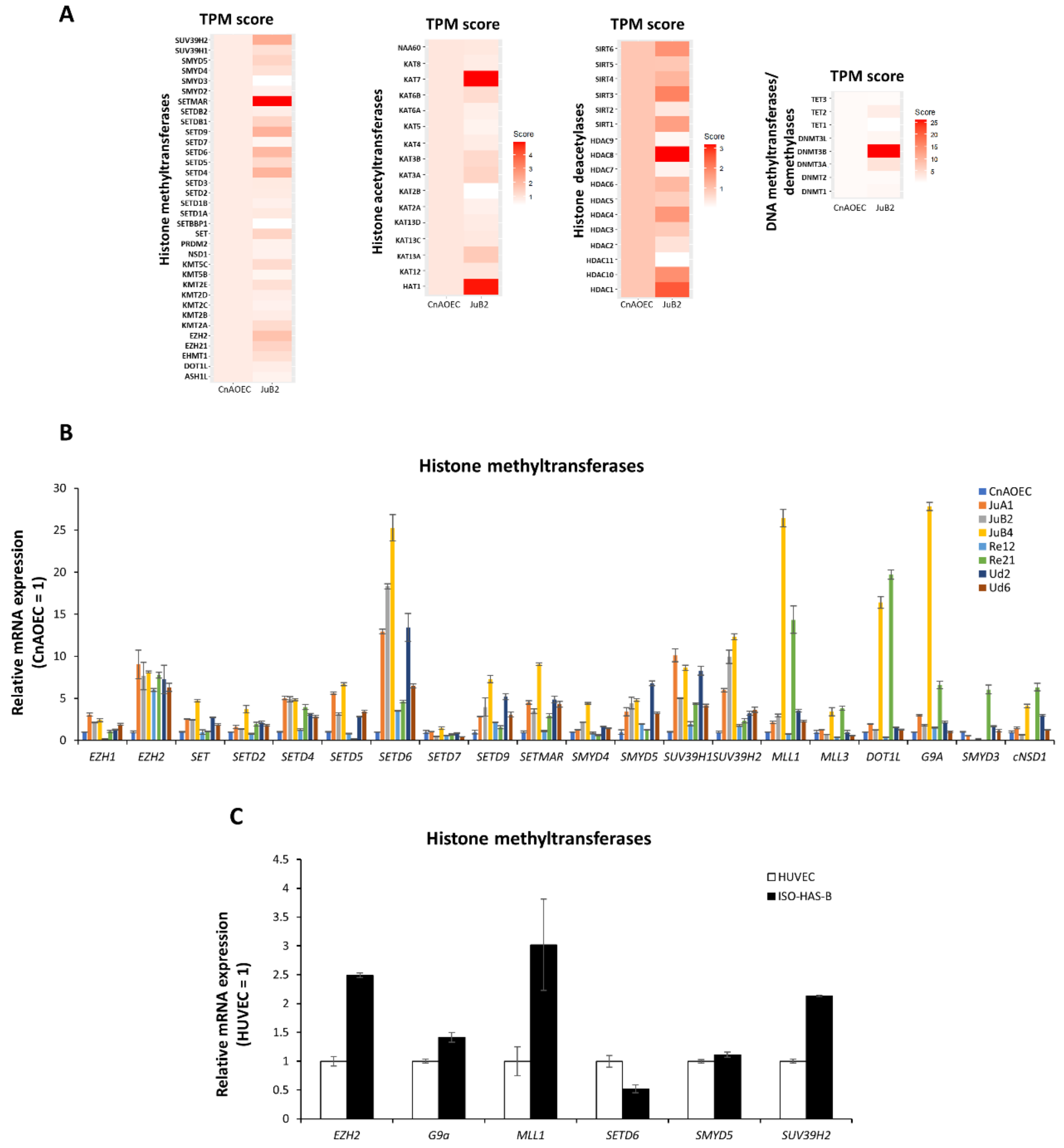


**Supplementary Fig. 1 Histone methyltransferases are differentially expressed in endothelial cell tumors.**

**A,** TPM scores of histone methyltransferases, histone acetyltransferases, histone deacetylases, DNA methyltransferases, and DNA demethylases in CnAOEC and JuB2 HSA cell line. **B,** qRT-PCR verification of select histone methyltransferases in CnAOEC and HSA cell lines. **C,** qRT-PCR verification of select histone methyltransferase in HUVEC and ISO-HAS-B cell lines. Data are presented as mean values ± s.d.

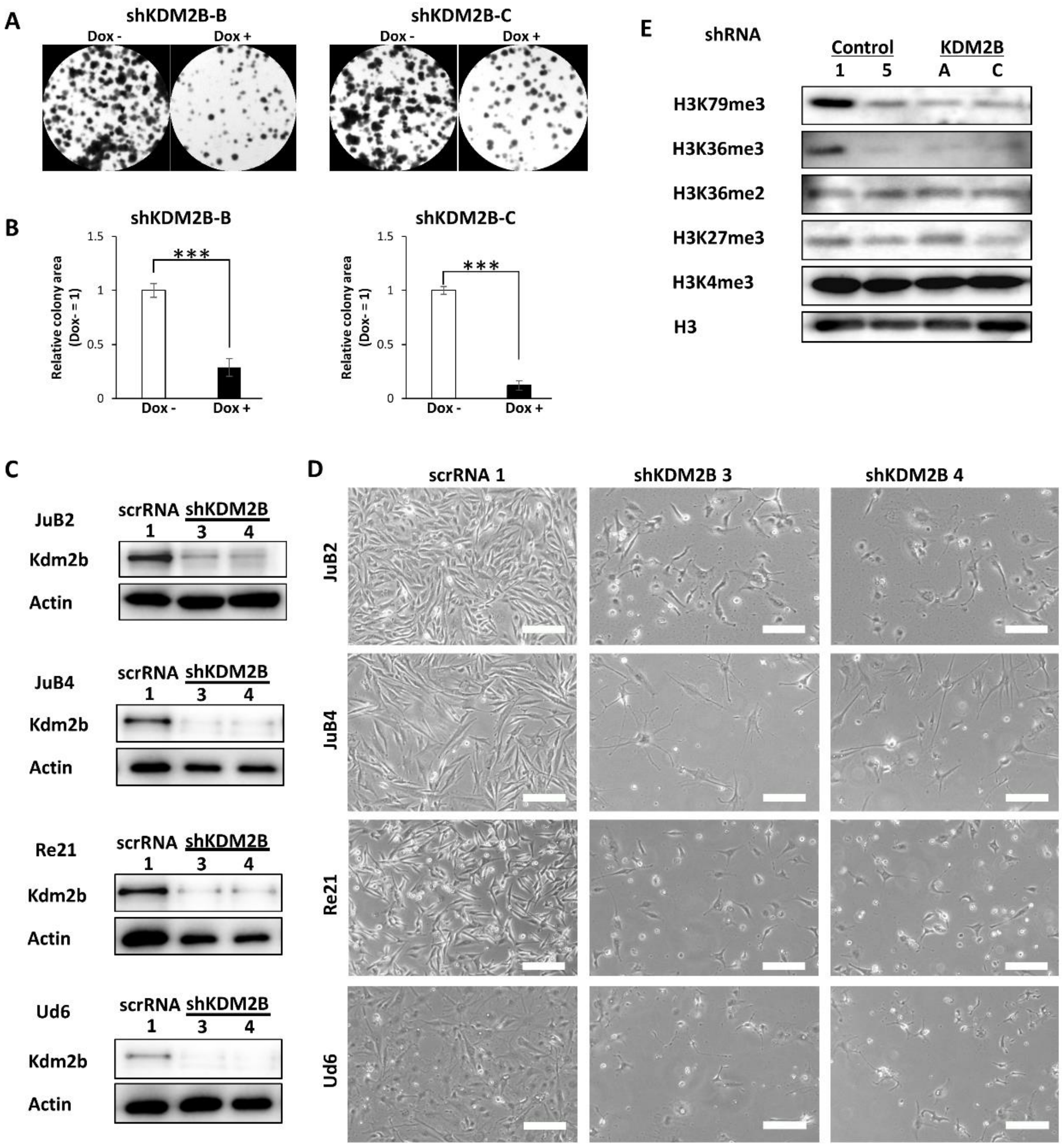


**Supplementary Fig. 2 Kdm2b is not a global demethylase in HSA.**

**A,** Colony formation assay of JuB2 cells after silencing of Kdm2b using shKdm2b-B and -C. **B,** Quantitative analysis of **A**. **C,** Western blotting for Kdm2b following Kdm2b knockdown with constitutively expressed silencing vectors. **D,** Viability and morphology of JuB2, JuB4, Re21, and Ud6 HSA cell lines 4 days after silencing of Kdm2b. Scale = 100 μm. **E,** Western blotting for histone lysine methylations after Kdm2b silencing. Data are presented as mean values ± s.d. ****p*<0.001, Student's *t*-test.

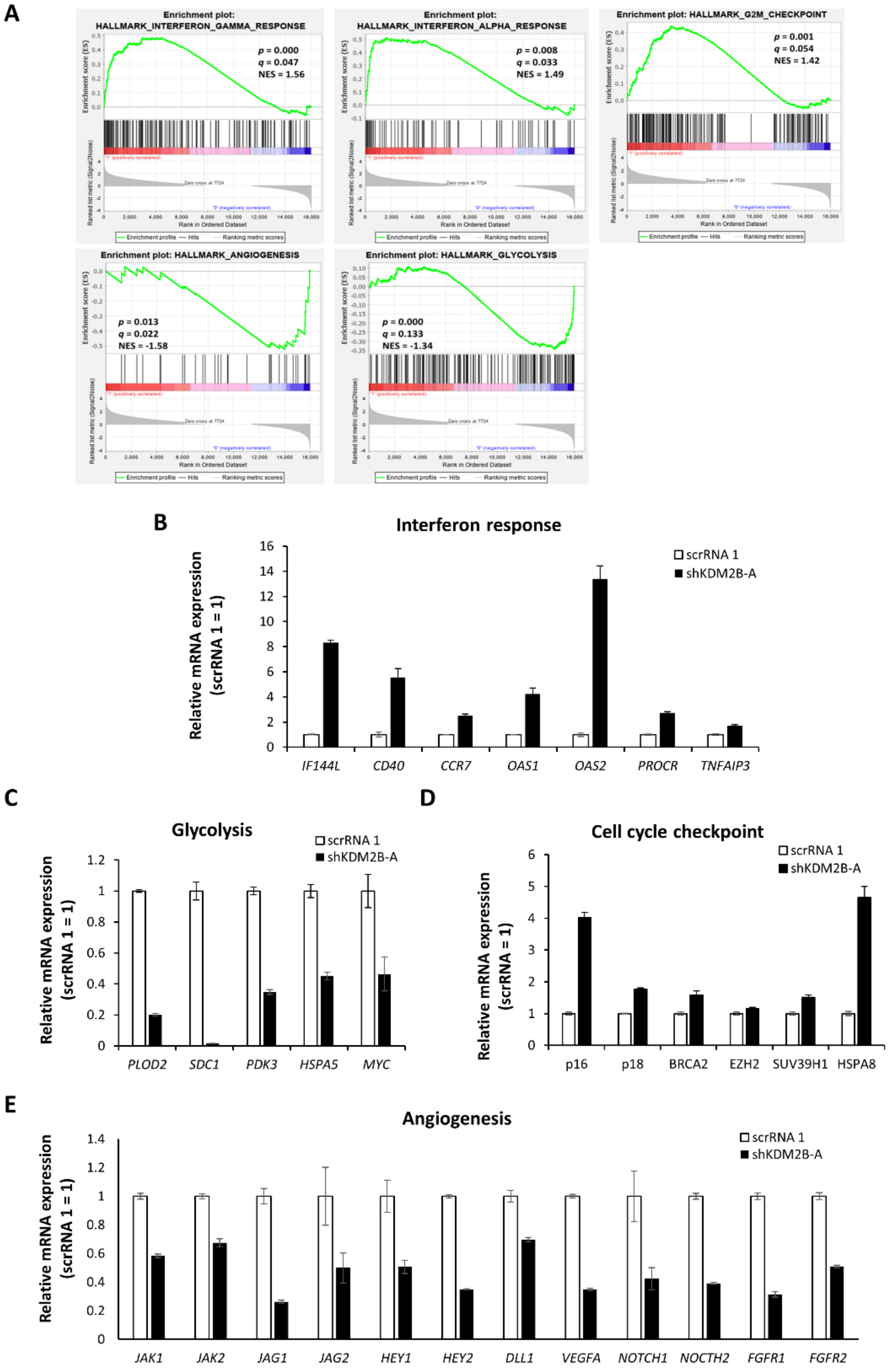


**Supplementary Fig. 3 Kdm2b regulates the interferon responses, cell cycle checkpoints, angiogenesis, and glycolysis pathways.**

**A,** GSEA enrichment plots in shKDM2B-A JuB2 cells versus scrRNA-1 JuB2 cells. **B to F,** qRT-PCR results of genes related to the enriched pathways in a. Data are presented as mean values ± s.d.

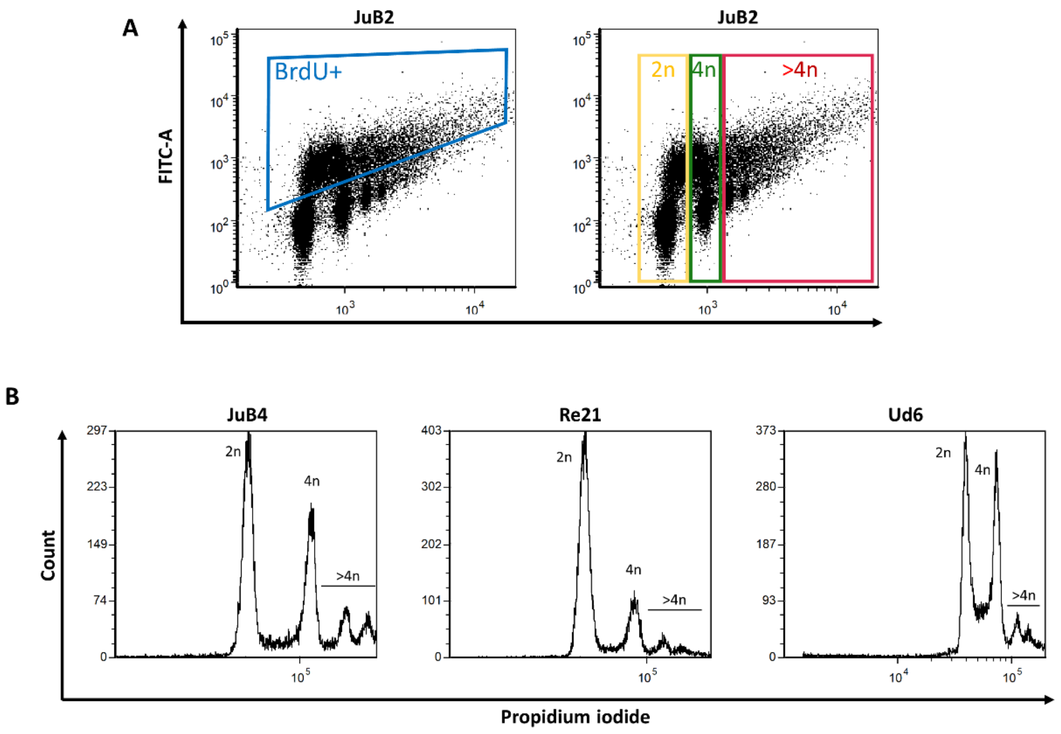


**Supplementary Fig. 4 Parental HSA cells are aneuploid.**

**A,** Dot plots illustrating the gating methods for 2n, 4n, >4n, and dividing cell populations.

**B,** Histograms of propidium iodide intensities in JuB4, Re21, and Ud6 cell lines.

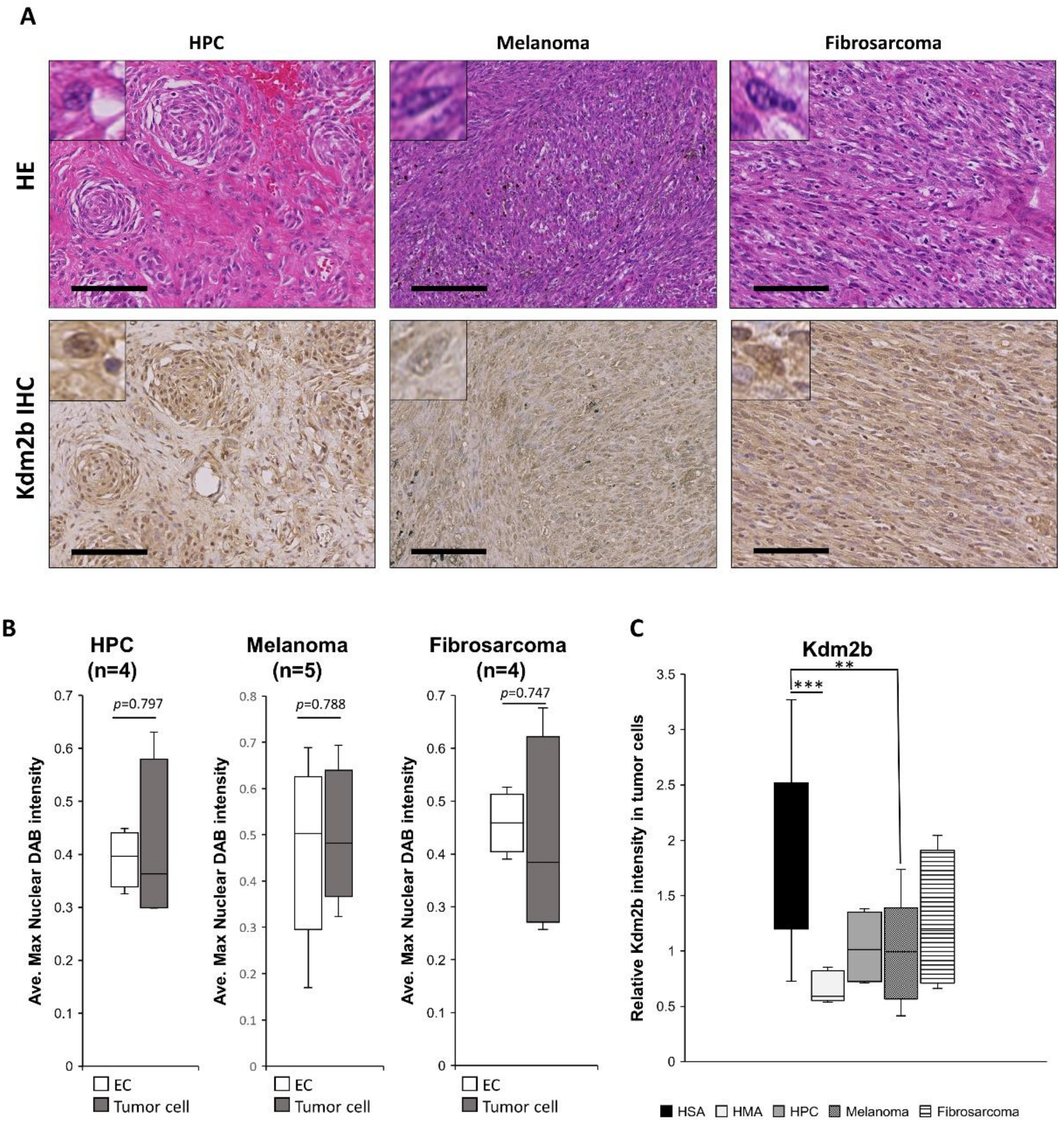


**Supplementary Fig. 5 Kdm2b expression can be used as a biomarker for endothelial cell tumors.** **A,** Representative images of HE and IHC analyses for clinical cases of HPC (n=4), melanoma (n=5), and fibrosarcoma (n=4). Scale = 100 μm. **B,** Average maximum nuclear DAB intensity in each HPC, melanoma, and fibrosarcoma case. **C,** Comparison of the relative Kdm2b intensities in tumor cells normalized by average Kdm2b intensities in EC on the same slides between HSA, HMA, HPC, melanoma, and fibrosarcoma. Data are presented as mean values ± s.d. ***p*<0.01, Bonferroni test.

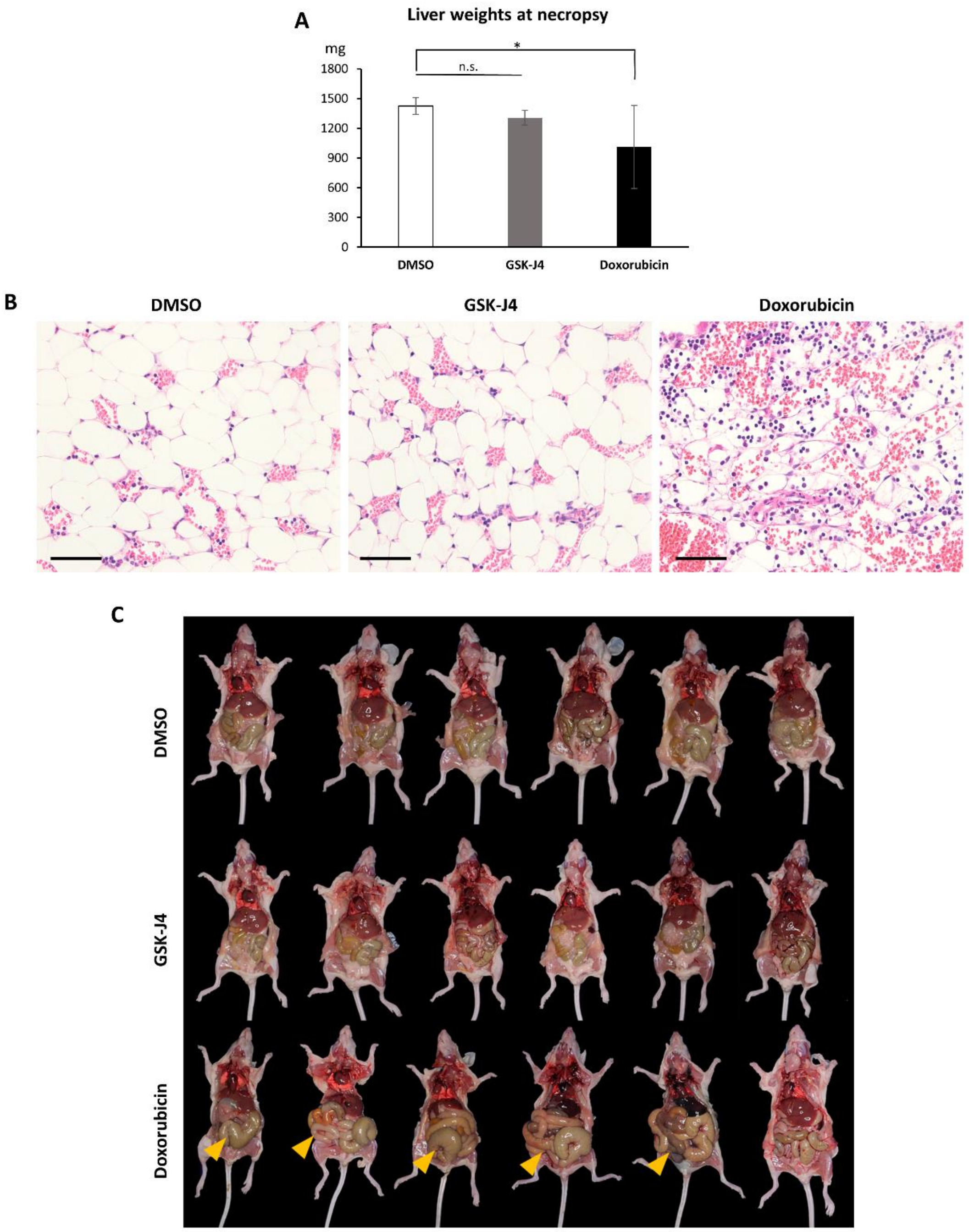


**Supplementary Fig. 6 GSK-J4 can be used as a therapeutic alternative to doxorubicin treatment in HSA.**

**A,** Liver weights of mice treated with DMSO, GSK-J4, or doxorubicin. Data are presented as mean values ± s.d. *$p$<0.05, Tukey's test. **B,** Representative images of bone marrow sections from mice treated wih DMSO, GSK-J4, or doxorubicin. Scale = 100 μm. **C,** Gross images of nude mice treated with DMSO, GSK-J4, or doxorubicin. Arrowheads indicate enlargement of intestinal segments.

**Supplementary Table 1**. List of antibodies used in the study.

| Protein Name | Catalog Number | Maker | Host | Dilution |
|---|---|---|---|---|
| Kdm1a | 2139S | Cell Signaling Technology | Rabbit | 1:1000 |
| Kdm2a | ab191387 | Abcam | Rabbit | 1:1000 |
| Kdm2b | sc-293279 | Santa Cruz Biotechnology, Inc. | Mouse | 1:1000 |
| Actin | MAB1501 | Sigma-Aldrich | Mouse | 1:1000 |
| p-Atm | sc-47739 | Santa Cruz Biotechnology, Inc. | Mouse | 1:1000 |
| t-Atm | sc-377293 | Santa Cruz Biotechnology, Inc. | Mouse | 1:1000 |
| c-Fos | sc-166940 | Santa Cruz Biotechnology, Inc. | Mouse | 1:1000 |
| γH2A.X | A300-081A-T | Bethyl Laboratories, Inc. | Rabbit | 1:1000 |
| Cleaved-caspase 3 | 9661S | Cell Signaling Technology | Rabbit | 1:1000 |
| p-Erk1/2 | 4370S | Cell Signaling Technology | Rabbit | 1:1000 |
| t-Erk1/2 | 4695S | Cell Signaling Technology | Rabbit | 1:1000 |
| H3K79me3 | 74073S | Cell Signaling Technology | Rabbit | 1:1000 |
| H3K36me3 | ab9050 | Abcam | Rabbit | 1:1000 |
| H3K36me2 | 39255 | Active Motif | Rabbit | 1:4000 |
| H3K27me3 | ab6002 | Abcam | Mouse | 1:1000 |
| H3 | 39140 | Active Motif | Rabbit | 1:5000 |
| Antirabbit | NA934VS | Sigma-Aldrich | Donkey | 1:10000 |
| Antimouse | G21040 | Thermo Fisher Scientific | Goat | 1:10000 |

**Supplementary Table 2**. List of qRT-PCR primers used in the study.

| Species | *Target* | Sequence | | Gene ID | Reference |
|---|---|---|---|---|---|
| | *TBP* | F | ATAAGAGAGCCCCGAACCAC | ENSCAFG00000004119 | Peters *et al.*, 2007 |
| | | R | TTCACATCACAGCTCCCCAC | | |
| | *KDM1A* | F | CCCACCTGAGGAAGAAAATG | ENSCAFG00000013361 | This paper |
| | | R | GGAAAACAGGCTGCTTCTTG | | |
| | *KDM2A* | F | AAGAAATGTGTCCCCACAGG | ENSCAFG00000011792 | This paper |
| | | R | CAAGCTCCTCCAGCAAAATC | | |
| | *KDM2B* | F | AGAAGCCACGAATGCCATTG | ENSCAFG00000008259 | This paper |
| | | R | AACAGCCTTTCACGCTCATC | | |
| | *KDM3A* | F | ACTTTAGGATGCCGGTCACAG | ENSCAFG00000007522 | This paper |
| | | R | TGCAGCTTCTTGAGTTTGGC | | |
| | *KDM4A* | F | GATTTCCCCTTTGATGCTGA | ENSCAFG00000004903 | This paper |
| | | R | TGGTAGACTCCGCACAGTTG | | |
| | *KDM4B* | F | CAGTCAGGCCTCTTCACACA | ENSCAFG00000018925 | This paper |
| | | R | CCAATACTTGCGTTCCAGGT | | |
| | *KDM4C* | F | CCACTGACTCTGGTGAAGCA | ENSCAFG00000001434 | This paper |
| | | R | TCGCCCAAGACTCTGTTTCT | | |
| e | *KDM5A* | F | GGACCTTGAGCCTCTGAGTG | ENSCAFG00000015781 | This paper |
| | | R | GGAATGCATGGCTTCAATCT | | |
| | *KDM5B* | F | GCTGTCCAACTCCCAAATGT | ENSCAFG00000010452 | This paper |
| | | R | GGCTCTTGGGTTTTTCCTTC | | |
| | *KDM6A* | F | ATAACCGCACAAACCTGACC | ENSCAFG00000014589 | This paper |
| | | R | AGGACCTGCCAAATGTGAAC | | |
| | *KDM6B* | F | TCTTCGATTTTCCCCCTACC | ENSCAFG00000016817 | This paper |
| | | R | GAATGGATTCGTCCAGCATC | | |
| | *KDM7A* | F | AGCAACCAGGCAACAAAAGG | ENSCAFG00000003990 | This paper |
| | | R | TCTTCCCAAGACGCTGTTTG | | |
| | *COL1A2* | F | TCTCCCTGGTGAATTTGGTC | ENSCAFG00000002069 | This paper |
| | | R | GTTCACCCTTGTTTCCATCG | | |
| | *INSIG1* | F | TGGGATCACTATTGCCTTCC | ENSCAFG00000005081 | This paper |
| | | R | AGCGGATGTAGAGAAAGTCTGG | | |
| | *ITGB3* | F | GGATTCCAGCAATGTCCTTC | ENSCAFG00000013735 | This paper |
| | | R | TGGCGTTGAACGATAGAGAC | | |
| | *PPARG* | F | TTCTCCAGCATTTCCACTCC | ENSCAFG00000004991 | This paper |
| | | R | AGGCTCCACTTTGATTGCAC | | |

| | | | | | |
|---|---|---|---|---|---|
| | *TGFBR3* | F | CACATTGTGCACCAAGAAGG | ENSCAFG00000020179 | This paper |
| | | R | TCATTGAGGCATCCAGTGAG | | |
| | *VLDLR* | F | GACGAACCCCTGAAAGAATG | ENSCAFG00000002028 | This paper |
| | | R | TGCGCAGTCACATTCATAGC | | |
| | *EZH1* | F | TGAGGAGTCCCTTTTTCGAG | ENSCAFG00000014827 | This paper |
| | | R | TCATCTGTTGGCAGCTTCAG | | |
| | *EZH2* | F | CAGACCGGTGAAGAGCTGTT | ENSCAFG00000003411 | This paper |
| | | R | GGGGAGGAAGAGGTAGCAGA | | |
| | *SET* | F | TCCATCGTCAAAGTCCACTG | ENSCAFG00000032728 | This paper |
| | | R | TCCTGCTGGCTTTATTCTGC | | |
| | *SETD2* | F | ACAGCAGAAGCAGACACCTC | ENSCAFG00000013392 | This paper |
| | | R | AGGCACTGGACGATGAACTG | | |
| | *SETD4* | F | TGAATCATAGCCCCGAAGTC | ENSCAFG00000009616 | This paper |
| | | R | AGCCGTTGGTTATCATGAGG | | |
| | *SETD5* | F | ACCCCCAAACACTACATTCG | ENSCAFG00000005486 | This paper |
| | | R | CCAAGGCTTGCTTTATCCAG | | |
| | *SETD6* | F | CAAACTCCCCTTTGATGGTG | ENSCAFG00000008461 | This paper |
| | | R | TTTAGGAATGGGCTGAGTGG | | |
| | *SETD7* | F | AGGTAGCGGTGGGACCTAAT | ENSCAFG00000003703 | This paper |
| | | R | GTTGTAGGGCTCAGGCACAT | | |
| | *SETD9* | F | AAGGCGCGGTTGTATCTATG | ENSCAFG00000006922 | This paper |
| | | R | TCAATGAGTACCCCATCCAG | | |
| | *SETMAR* | F | CTTGGAGAACGTGCCTGTGA | ENSCAFG00000005958 | This paper |
| | | R | CAAATGCATCCGGGAAAGGT | | |
| | *SMYD3* | F | CGGAGATGCAGGAAGTTGGT | ENSCAFG00000029171 | This paper |
| | | R | CTCGATGTCTCGGACTGCTC | | |
| | *SMYD4* | F | TGTGGGAAAGGACCCTAATG | ENSCAFG00000019244 | This paper |
| | | R | AGTGTGAGGCCTTGAATGTG | | |
| | *SMYD5* | F | TGAAGGATCTGGCCTGTATG | ENSCAFG00000008932 | This paper |
| | | R | TGGCTTGATATCCTCCAAGG | | |
| | *SUV39H1* | F | TGGAGAAGATCCGCAAGAAC | ENSCAFG00000015545 | This paper |
| | | R | TGTACACGTCCTCCACGTAGTC | | |
| | *SUV39H2* | F | CGATTGGAATCACCAAAAGG | ENSCAFG00000004696 | This paper |
| | | R | AGAATCTGGCCATCCTTTCC | | |
| | *MLL1* | F | GCTTTGGCTCCAGCAAGAAC | ENSCAFG00000012691 | This paper |
| | | R | CGTCAGTGACTTCCAGGCAT | | |

| | | | | | |
|---|---|---|---|---|---|
| | *MLL3* | F | TCGCTCCAAGAAAAGGAAGA | ENSCAFG00000004955 | This paper |
| | | R | CAAGCCATAGGAGGTGGTGT | | |
| | *DOT1L* | F | ACCACGATGCTGCTCATGAA | ENSCAFG00000019420 | This paper |
| | | R | GCCTCTGCATGCTTTCGAAG | | |
| | *G9A* | F | GAAGAAGTGGCGGAAGGACA | ENSCAFG00000000669 | This paper |
| | | R | ACTCACTAGGGCCTGAGGAG | | |
| | *NSD1* | F | TATGGAGGGGGATGTGAGCA | ENSCAFG00000016473 | This paper |
| | | R | GGTCTTCCTGCAGCTGTCTT | | |
| | *P15INK4B* | F | GTGCGGCAGCTCCTGGAAGC | ENSCAFG00000001675 | This paper |
| | | R | GCCCATCATCATGACCTGGATCG | | |
| | *P16INK4A* | F | GTGGACCTGGCTGAGGAGCG | ENSCAFG00040004457 | This paper |
| | | R | TTCTTGAAGTCCGGGCTGTCTG | | |
| | *ATR* | F | CAGCGCTTCCTAGTACTCCG | ENSCAFG00000007863 | This paper |
| | | R | TTGGCAGCAAGGTCAGGTAG | | |
| | *IF144L* | F | TCTATTTTCCGAGGCCAGAG | ENSCAFG00000020343 | This paper |
| | | R | TCATGTATCCCCATGGAGTC | | |
| | *CD40* | F | TATTCACCTCGCCATGGTTC | ENSCAFG00000009994 | This paper |
| | | R | GAGTGCATTCCGTGTCAATG | | |
| | *CCR7* | F | GGCTCTCCTTGTCATTTTCC | ENSCAFG00000030300 | This paper |
| | | R | TCCACCGTGGTATTTTCTCC | | |
| | *OAS1* | F | TGTGCGGGTGTCTAAAGTTG | ENSCAFG00000023556 | This paper |
| | | R | TGAACTGTCCTCGTTTCTCG | | |
| | *OAS2* | F | TGACCCAGATCCAGAAAACC | ENSCAFG00000023107 | This paper |
| | | R | CCATTCGGTAGCGTCTTTTG | | |
| | *PROCR* | F | GCAGGAACACAATGCTTCAA | ENSCAFG00000007945 | Aoshima *et al.*, 2018 |
| | | R | AAGATGCCTACAGCCACACC | | |
| | *TNFAIP3* | F | GGTGATCGAAATTCCTGTCC | ENSCAFG00000000267 | This paper |
| | | R | TGGGTAAGTTGGCTTCATCC | | |
| | *BAX* | F | ACATGGAGTTGCAGAGGATG | ENSCAFG00000003867 | This paper |
| | | R | CCAGTTGAAGTTGCCATCAG | | |
| | *RB1* | F | AAGCAGAAGCCAACTTGACCAG | ENSCAFG00000004436 | This paper |
| | | R | GTCCTTCTCGGTCTTTTGCTTG | | |
| | *MCL1* | F | AGCTGCATCGAACCATTAGC | ENSCAFG00000012050 | This paper |
| | | R | AGAACTCCACAAACCCATCC | | |
| | *E2F1* | F | GATGGTCATGGTGATCAAGG | ENSCAFG00000007429 | This paper |

| | | | | | |
|---|---|---|---|---|---|
| | | R | GCACAGGAAAACGTCAATGG | | |
| | *CDC25A* | F | CCTGAAAAGGAGCCATTCTG | ENSCAFG00000012919 | This paper |
| | | R | ACGGGGTCTCTTCATCATTG | | |
| | *p18* | F | GCTGCAGGTTATGAAACTTGG | ENSCAFG00000028905 | This paper |
| | | R | CGAAACCAGTTCGGTCTTTC | | |
| | *BRCA2* | F | CGGGAGATTGACTGTGGGTC | ENSCAFG00000006383 | This paper |
| | | R | GCAAGCAGGACGAGTACTGT | | |
| | *HSPA8* | F | AACCACCCCAAGTTATGTCG | ENSCAFG00000011666 | This paper |
| | | R | TCAGATTGGACGACAGCATC | | |
| | *JAK1* | F | TACGGACAACATCAGCTTCG | ENSCAFG00000018615 | This paper |
| | | R | TGAGGATCCGGTCAAAACTC | | |
| | *JAK2* | F | TCAGATGTCTGGAGCTTTGG | ENSCAFG00000002102 | This paper |
| | | R | TCATCTGTCCTTGCTTGTCG | | |
| | *JAG1* | F | GAAGCGTGGGATTCCAGTAA | ENSCAFG00000005627 | This paper |
| | | R | CAGAACTTGTTGCAGCCAAA | | |
| | *JAG2* | F | AGGGCAGTACCTGCAACATC | ENSCAFG00000018401 | This paper |
| | | R | TGCAGGAGAACGAGTCAATG | | |
| | *HEY1* | F | GCGCGGATGAGAATGGAAAC | ENSCAFG00000008391 | Aoshima *et al.*, 2018 |
| | | R | GTCGGCGCTTCTCAATGATG | | |
| | *HEY2* | F | CGGCGAGATCGGATAAATAA | ENSCAFG00000032212 | Aoshima *et al.*, 2018 |
| | | R | CGCGTCGAAGTAGCCTTTAC | | |
| | *DLL1* | F | CCGATGACCTCACAACAGAA | ENSCAFG00000004094 | This paper |
| | | R | GCAGACCTTCTCCCCTCTCT | | |
| | *VEGFA* | F | TATGGCAGGAGGAGAGCACAAACC | ENSCAFG00000001938 | This paper |
| | | R | CAGCCCCCACACCGCATCAG | | |
| | *NOTCH1* | F | TACCGGCCAGAACTGTGAGGAGAA | ENSCAFG00000019633 | Aoshima *et al.*, 2018 |
| | | R | GGAGGGCAGCGGCAGTTGTAAGTA | | |
| | *NOCTH2* | F | TCGGGATAGCTATGAGCCCT | ENSCAFG00000010476 | Aoshima *et al.*, 2018 |
| | | R | GGCATGTTGCTTTCCCCAAC | | |
| | *FGFR1* | F | TCCGTCAATGTCTCAGATGC | ENSCAFG00000005970 | This paper |
| | | R | CCATCTTTTCTGGGGATGTC | | |
| | *FGFR2* | F | TCCGTCAATGTCTCAGATGC | ENSCAFG00000012374 | This paper |
| | | R | CCATCTTTTCTGGGGATGTC | | |

| | | | | | |
|---|---|---|---|---|---|
| | *PLOD2* | F | TATGGCTCTCTGCCGAAATG | ENSCAFG00000008101 | This paper |
| | | R | TGGAATGTTTCCGGAGTAGG | | |
| | *SDC1* | F | TCTGGGGATGACTCTGACAAC | ENSCAFG00000003833 | This paper |
| | | R | TCTGCTGTGACAAGGTGATG | | |
| | *HSPA5* | F | CGGAGGCTTATTTGGGAAAG | ENSCAFG00000020196 | This paper |
| | | R | TGGGCATCATTGAAGTAGGC | | |
| | *PDK3* | F | TGTCCATCAAGCAGTTCCTG | ENSCAFG00000013476 | This paper |
| | | R | AGCAAGTTATCCGGCAGAAG | | |
| | *MYC* | F | CGCTGGTCCTTAAGAGATGC | ENSCAFG00000001086 | This paper |
| | | R | CGCCTCTTGTCATTCTCCTC | | |
| Human | *KDM1A* | F | CCCATGGAAACTGGAATAGC | ENSG00000004487 | This paper |
| | | R | GCCAAGCTTTCATCCATCTC | | |
| | *KDM2A* | F | AAGAAATGTGTCCCCACAGG | ENSG00000173120 | This paper |
| | | R | CAAGCTCCTCCAGCAAAATC | | |
| | *KDM2B* | F | ATGCCTGACCCTGATTTCAC | ENSG00000089094 | This paper |
| | | R | TGGGTGTTCACATCCATCAC | | |
| | *EZH2* | F | TTTCCAGATAAGGGCACAGC | ENSG00000106462 | This paper |
| | | R | ATGTTGGGGGTACATTCAGG | | |
| | *G9A* | F | GGTTTGCGCTTCAACTCAAC | ENSG00000204371 | This paper |
| | | R | AATGGGCACGTTCTCATAGC | | |
| | *MLL1* | F | TGCCTGGAAGTCATTGACAG | ENSG00000118058 | This paper |
| | | R | GGAACACAACTGCATCATGG | | |
| | *SETD6* | F | AGGATGAAAAGGAGCCCAAC | ENSG00000103037 | This paper |
| | | R | TTTAGGAATGGGCTGAGTGG | | |
| | *SMYD5* | F | TGAAGGATCTGGCCTCTTTG | ENSG00000135632 | This paper |
| | | R | AAAGGAGGTCTCTGCATTGG | | |
| | *SUV39H2* | F | TCGATACGGCAATGTGTCTC | ENSG00000152455 | This paper |
| | | R | CAATGCTATTCGGGGAAGAC | | |

**Supplementary Table 3**. List of shRNA oligos used in the study.

| Species | Target | Sequence | Reference |
|---|---|---|---|
| Canine | KDM1A-A | AAGUGAUACUGUGCUAGUCCAC | This paper |
| | KDM1A-B | ACUUCAGGAUGUGAAGUGAUAG | This paper |
| | KDM1A-C | GUAUAGAGCAAGAGAAGCAGAU | This paper |
| | KDM2A-A | AGCUCUCAGUGGCAUCAUCAAG | This paper |
| | KDM2A-B | CAUGCUGUAUCUGCAAUGAGAU | This paper |
| | KDM2A-C | CGCCCACAACCUGGAGCUGUAC | This paper |
| | KDM2B-A | ACCACAGCAUCUGAAGGAGAAG | This paper |
| | KDM2B-B | CCCAGUGCCUGUCCUUCUUCAA | This paper |
| | KDM2B-C | CCCAGAGAGAAUCCAUGCUUAU | This paper |
| | KDM2B-3 | UGGAAAUAUCUGUCAUAUUGA | This paper |
| | KDM2B-4 | CAGUUCAUAGCUGAAAUGUCU | This paper |
| | scramble 1 | AUAAGAGACGAACGUAACAUA | This paper |
| | scramble 5 | AGAAGAUUAACGUAGAAGGUG | This paper |

**Supplementary Table 4**. List of oligos used for the induction of mutations in KDM2B sequence.

| Species | Target | Sequence | | Reference |
|---|---|---|---|---|
| Canine | WT Kdm2b | F | CGCTACCGGTCTCGAGACCATGCATCGGGCAGTGGACCCTC | This paper |
| | | R | CGACGGTACCGAATTCCAGAGGCGGGACCTAGGTCCAGC | |
| | Silent mutation for shKdm2b C | F | CGCGAGTCTATGCTGATTGATGCCCCAAGAAAGCC | This paper |
| | | R | CAGCATAGACTCGCGTTGGTATTCCTGAGTGAGGT | |
| | $Kdm2b^{H283Y}$ | F | TGGATTTACGCAGTCTACACCCCCGTA | This paper |
| | | R | GACTGCGTAAATCCAACCGGAAGG | |
| | $Kdm2b^{C587A}$ | F | ACGCGAGCCCGCAAGTGCGAGGCC | This paper |
| | | R | CGCCGGACGCGAGCCCGCAAG | |
| | $Kdm2b^{\triangle PHD}$ | F | GCGCCAGTGCTGCCCCATGGCAAGACCGGGAAACAAAA | This paper |
| | | R | TTTCCCGGTCTTGCCATGGGGCAGCACTGGCGCGA | |
